# Kinetic asymmetry drives directionality in an ATP-binding cassette transporter

**DOI:** 10.64898/2026.09.11.751009

**Authors:** Michael Rudolph, Hossein Batebi, Malavika Pramod, Katja Barth, Clemens Hirschberg, Robert Tampé, Roland R. Netz, Benesh Joseph

**Author notes:** These authors contributed equally (Michael Rudolph, Hossein Batebi).

## Abstract

How ATP binding and hydrolysis directionally reshape the conformational landscape remains unknown for ATP-binding cassette (ABC) transporters. Here, we identify two conserved ionic locks within the nucleotide-binding domains that govern transition barriers and energy transduction: an intra-subunit inward-facing (IF)-lock and an inter-subunit outward-facing (OF)-lock. Mg^2+^-ATP acts as a molecular key that disrupts the IF-lock, driving the forward transition. Following ATP hydrolysis, release of the γ-phosphate, which, together with Mg^2+^, forms the pivot of the OF-lock, initiates the reverse transition. Directionality arises from kinetic asymmetry, driven by an anticorrelated exchange of the rate-limiting step between the consensus nucleotide-binding site and the transmembrane domains during forward and reverse transitions, respectively. Conservation of these molecular locks reveals a universal blueprint for ATP-driven mechanical transduction across the ABC superfamily.

## Introduction

ABC transporters are among the largest and most widespread protein superfamilies (*1-3*). Their dysfunction is associated with numerous human diseases, and they contribute to multidrug resistance (*2, 4, 5*). ATP binding and hydrolysis at the nucleotide-binding domains (NBDs) are transmitted to the transmembrane domains (TMDs), which create a pathway for substrate permeation across the membrane. Despite extensive structural and computational studies, the molecular determinants that define the transition barriers of the transport cycle, and how the measured kinetics arise from them, have remained unknown. Furthermore, the physical principles dictating how such systems achieve directionality remain poorly understood. A mechanistic understanding of energy transduction therefore requires quantitative mapping of conformational equilibria with sufficient spatiotemporal resolution (*6-11*). Here, we address these challenges for the ABC exporter TmrAB from *Thermus thermophilus*. By combining time-resolved (TR) electron–electron double resonance (PELDOR/DEER) spectroscopy (*12-14*), unbiased molecular dynamics simulations (*15-20*), and non-Markovian rate theory (*21-23*), we successfully resolve both the conformational heterogeneity (*24-28*) and the underlying kinetics. We show that two conserved ionic locks define the activation barriers and mechanically transduce ATP binding and hydrolysis into directional conformational changes and substrate translocation.

## Results

TmrAB remains active over a broad temperature range (30 – 70 °C), and high-resolution structures are available for multiple conformational states (*29, 30*). It is a heterodimer containing one ATPase-active consensus nucleotide-binding site (c-NBS) and one catalytically impaired degenerate nucleotide-binding site (d-NBS) (*31*). We engineered spin pairs at the two NBSs and the TMDs in a cys-less background (denoted as the parental strain) and quantified their closed ↔ open equilibrium using TR-PELDOR (**Figures 1a–c**). All these spin-labeled variants actively hydrolyzed ATP (*14*). Globally, the transport cycle in TmrAB primarily involves three major conformations, namely, inward-facing (IF), occluded (OCC), and outward-facing (OF) states (**Figure 1a**), which might be connected via transient intermediate states (*30*).

**Figure 1.**
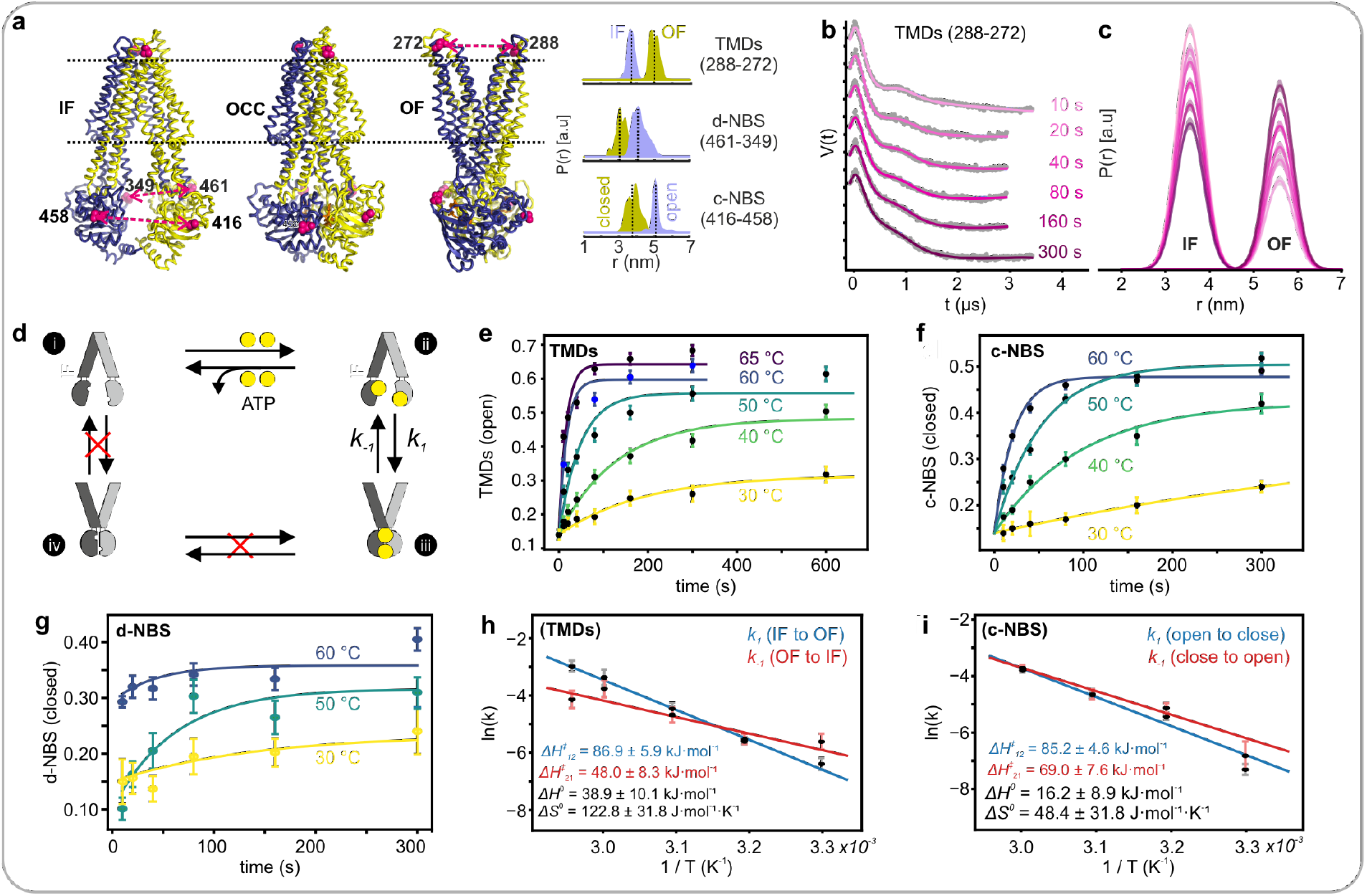
Kinetics and thermodynamics of IF ↔ OF transition in TmrAB. (**a**) The IF (PDB ID: 6RAG), OCC (PDB ID: 6RAI), and OF (PDB ID: 6RAH) structures are shown with spin-labeled positions on the TMDs (288-272), c-NBS (416-458), and the d-NBS (461-349) highlighted as spheres. Simulations for the inter-spin distances are presented on the right. **(b–c**) A typical TR-PELDOR data set (50 °C) and the determined IF ↔ OF equilibrium for the TMDs are shown (Supplementary figures 1-3). (**d**) Kinetic model for the IF ↔ OF transition. As ATP is required for the transition and ATP cannot bind to closed NBSs, i ↔ iv and iii ↔ iv transitions are excluded. *k*_*1*_ and *k*_*-1*_ correspond to a two-state transition for the TMDs (closed ↔ open) or the NBDs (open ↔ closed), respectively. (**e–g**) Kinetics of IF ↔ OF transition as determined from TR-PELDOR. Sampling was mostly limited to 10–300 s to exclude deviations due to prolonged incubation. The fits from the kinetic model (corresponding to steps ii ↔ iii in panel d) are overlaid. Error bars indicate 95% confidence intervals. (**h-i**) Analysis of the rate constants (as determined from e–f, with the SE values shown) according to Kramers’ reaction rate theory for the forward (blue) and reverse (red) transitions.

Initially, we observed the IF-to-OF transition under non-hydrolyzing conditions in the presence of saturating ATP (+EDTA). Under this condition, the kinetic model simplifies to the transition between ATP-bound IF and OF conformations (**Figure 1d**). In agreement, the TR-PELDOR data gave an excellent fit in the global analysis (*32*) as a combination of open and closed states (**Figure 1b–c, Supplementary figures 1–3**). Further, the determined populations well fitted into a single-exponential (first-order) relaxation to equilibrium model (step **ii** ↔ **iii** in **Figure 1d, Supplementary Tables 1** and **2**). The d-NBS exhibited significantly faster kinetics, preventing sufficient resolution for reliable analysis (**Figure 1g, Supplementary Figure 3**). It maintains a smaller closed population, implying inherent flexibility independent of the TMDs (as further supported by MD simulations) under the non-hydrolyzing conditions.

Further analysis of the rate constants according to Kramers’ reaction rate theory (**Figures 1h–i**, blue lines) gave a similarly elevated activation enthalpy (Δ*H*^*‡*^) for the c-NBS and TMDs for the forward (IF → OF) transition (85.2 ± 4.6 and 86.9 ± 5.9 kJ·mol^-1^, respectively), in agreement with the thermophilicity of TmrAB. The reverse transition (OF → IF) at the c-NBS and the TMDs (**Figures 1h–i**, red lines) gave Δ*H*^*‡*^ values of 69.0 ± 7.6 and 48.0 ± 8.3 kJ·mol^-1^, respectively. This pronounced enthalpic divergence reveals a decoupled resetting mechanism, where the reverse transition is bottlenecked by a larger barrier at the c-NBS under non-hydrolyzing conditions.

Despite the large enthalpic factor for the IF → OF transition, a corresponding entropy gain offsets the Δ*G*^*0*^ ≈ 0 (**Figures 1h–I**, see methods) for both c-NBS and TMDs at 60 °C, revealing a similar position along the free energy landscape near physiological temperature. Experiments employing nanodisc-reconstituted TmrAB showed somewhat faster kinetics (**Supplementary figures 4a–e**). A 1:1 addition of the peptide substrate does not alter the (IF **↔** OF) equilibrium in TmrAB (*33*). Also, peptide transport shows a similar temperature dependence to the structural transition (**Supplementary figure 4f**), suggesting that the conformational changes themselves are the major limiting step for transport.

As closure of the c-NBS is a prerequisite, TMD opening might rapidly follow and therefore reflect the same apparent activation barrier as c-NBS. We resorted to unbiased molecular dynamics (MD) simulations to resolve the sequence of the conformational transitions (60 °C / 333.15 K) in POPE/POPG (3:1) membranes). In addition to the wild-type (WT), we also tested the E523Q (E→Q) variant in which the catalytic Glu (c-Glu) at the c-NBS is substituted with a Gln (*34*). For both variants, two of the ten replicas progressively approached the OCC state in the presence of Mg^2+^-ATP (**Supplementary figures 5–I, 5–II**). Principal-component analysis (PCA), combined with HDBSCAN clustering (*35*), revealed five trajectory centroids (C0 → C4, **Figure 2a, Supplementary figures 5–III**). The C0 – C4 clusters form sequential intermediates along the IF → OCC transition, while a separate trajectory connects the OCC and OF conformations. In line with the spectroscopic data (**Figure 1g**), the d-NBS contracts earliest and most strongly, dropping from ∼41 Å in C0 to ∼31 Å in C3. The c-NBS lags, showing closure only in later clusters. The intracellular loop (ICL, at the TMDs, **Supplementary figure 6f**) closure follows progressively (16 Å → 9 Å) towards the OCC state, paralleling the later stages of the c-NBS approach. We further initiated six independent (1.7 µs each) simulations from the occluded structure. One of the replicas briefly visited the OF conformation during the simulation time for the E → Q variant (**Figure 2a**, OCC → OF, **Supplementary figure 6**). In this run, the ICL and c-NBS remained closed (with the d-NBS exhibiting significant flexibility), indicating that TMD opening can occur after/with stable NBD engagement. Overall, these results strongly support an ATP-driven IF → OCC → OF transition with asymmetric timing: d-NBS closes first, c-NBS and ICL follow, and TMD opening as the last step.

**Figure 2.**
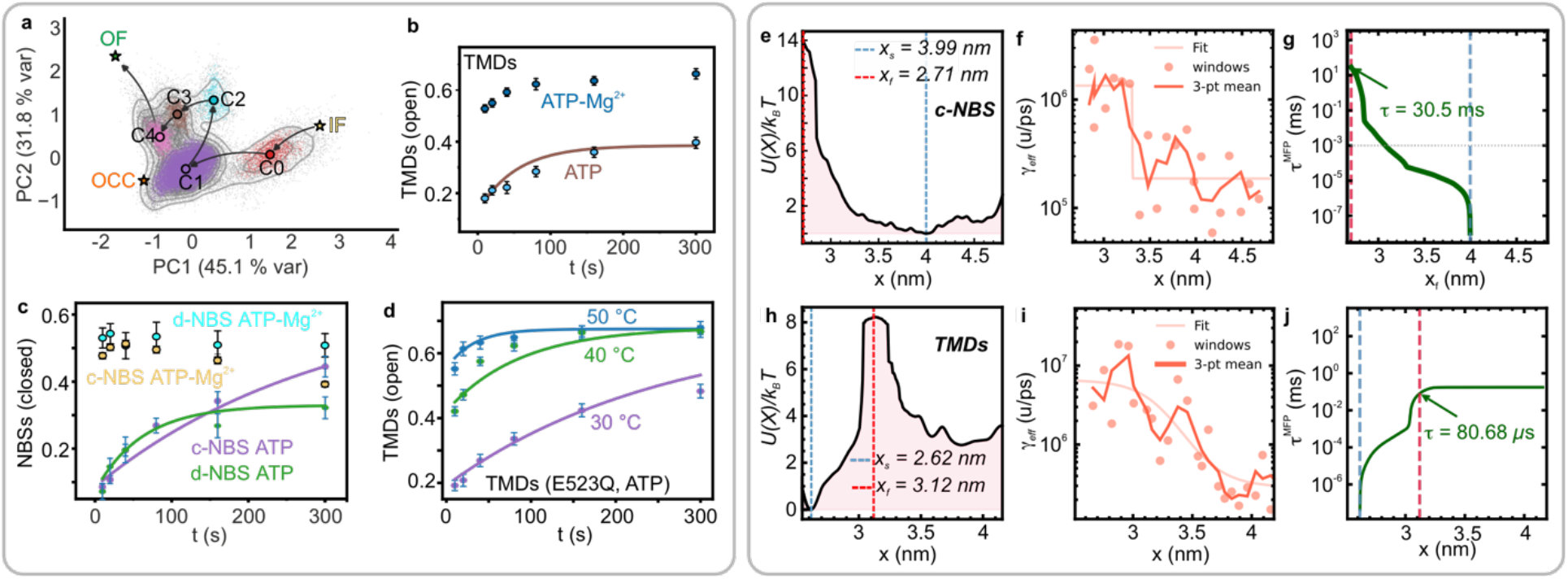
Molecular dynamics simulations, effect of Mg^2+^ ions, and MFPT calculations for the forward transition (IF → OF). (**a**) PCA analysis of 10 × 500 ns unbiased MD simulations for the IF → OCC → OF transition for the E→Q variant of TmrAB (**Supplementary figures 5**). Four C_α_–C_α_ distances (C416–L458 (c-NBS), D349–V461 (d-NBS), and V288-E272 (TMDs), as established with PELDOR spectroscopy, and D142–T124 (ICL, **Supplementary figure 6f**)) were z-scored and equally weighted (1/4 each) as the features for the PCA analysis. Points are colored by HDBSCAN cluster; gray dots are noise, and centroids Cn are ordered sequentially. Reference states are marked by stars. (**b**) Effect of ATP-Mg^2+^ on the IF ↔ OF transition at the TMDs, (**c**) c-NBS and the d-NBS (**Supplementary figures 7a–c)**. Data with ATP alone (marked as ATP) is overlaid for a direct comparison. Data were acquired at 40 °C, except that the reference d-NBS ATP samples (green curve in panel c) were observed at 50 °C. (**d**) Temperature-dependent IF ↔ OF transition observed at the TMDs in the E→Q variant in the presence of ATP (without Mg^2+^, **Supplementary figures 7d–f**). (**e-j**) MFPT analysis for the c-NBS and TMDs; the calculated free-energy (e, h), friction (*γ*_*eff*_, f, i), and MFPT profiles (g, j) are shown (see **Supplementary figures 8–17**).

In view of the accelerated transition observed from the MD simulations, we experimentally tested the effect of Mg^2+^ on the kinetics. Strikingly, Mg^2+^-ATP (vs. ATP) enhanced transitions at the two NBSs and the TMDs into the milliseconds range, even at 40 °C (313.15 K, **Figure 2b-c, Supplementary figures 7a–c**). A similarly accelerated transition was observed in the E→Q variant without Mg^2+^ (ATP alone, **Figure 2d, Supplementary figures 7d–f)**. To elucidate the rate-limiting step, we performed additional targeted MD simulations (at 310 K), starting with the Mg^2+^-ATP-bound IF structure of the WT protein, moving to OCC and OF conformations. Subsequently, umbrella sampling was used to calculate the transition barriers (**Figures 2e–j**, and **Supplementary figure 8**), friction profiles (γ_eff_) (*21, 22, 36, 37*) (**Figures 2e–j, Supplementary Figures 8–16**), and the mean first-passage times (MFPTs, **Figures 2g, j**) along the three collective variables (CVs), as established by the PELDOR experiments (c-NBS, d-NBS, and TMDs). The c-NBS closure revealed the slowest MFPT in the millisecond range (**Figure 2h**). In contrast, TMD opening (OCC → OF) gave an MFPT in the microsecond timescale (**Figure 2i**) and the d-NBS closed in the sub-microsecond (hundreds of nanoseconds) range (**Supplementary figure 8**). We computed the friction (*γ*) by three independent methods (*36, 38, 39*) on the same umbrella data. All three place c-NBS closure in the millisecond regime (*γ*_*eff*_ = 30.5 ms, *γ*_*pos*_ = 94.4 ms, and *γ*_*sol*_ = 324.7 ms), matching the TR-PELDOR kinetics (**Figures 2b–c)** and confirming it as the rate-limiting step of the IF → OF transition. Our memory-aware *γ*_*eff*_ gives the fastest time, consistent with memory-accelerated barrier crossing (*21, 22*) (**Supplementary figure 18**). Unlike the TMDs (**Figures 2h–j**), the MFPT profiles for both NBSs do not reach a plateau, suggesting a strained conformation.

Having localized the rate-limiting barrier, we next sought its structural origin. Based on the observations that Mg^2+^ or the E→Q substitution accelerates the IF → OF transition (**Figures 2b–d**), we further mapped the contacts surrounding the c-Glu as well as the interfacial interactions using MD simulations. These simulations revealed that in the IF conformation, the c-Glu (TmrA-E523), forms a salt bridge triad with the H-loop His and the nearby Arg (TmrA-H554 and TmrA-R555, respectively), thereby forming an intradomain ionic lock (named as IF-lock) in the NBD of TmrA (**Figure 3a, Supplementary figure 19)**. TmrA-R555 is further stabilized through an intradomain interaction with the conserved TmrA-Glu533, which is four amino acids away from the invariant D-loop TmrA-Asp529 residue. In TmrB, the same ionic constellation exists (**Figure 3a, Supplementary figure 20**). However, the salt bridges and the ionic lock are not intact due to suboptimal orientation between the residues, which might account for the lower barrier for d-NBS closure. Overall, a set of four conserved residues forms an intradomain ionic lock near each NBS, thereby stabilizing the IF conformation.

**Figure 3.**
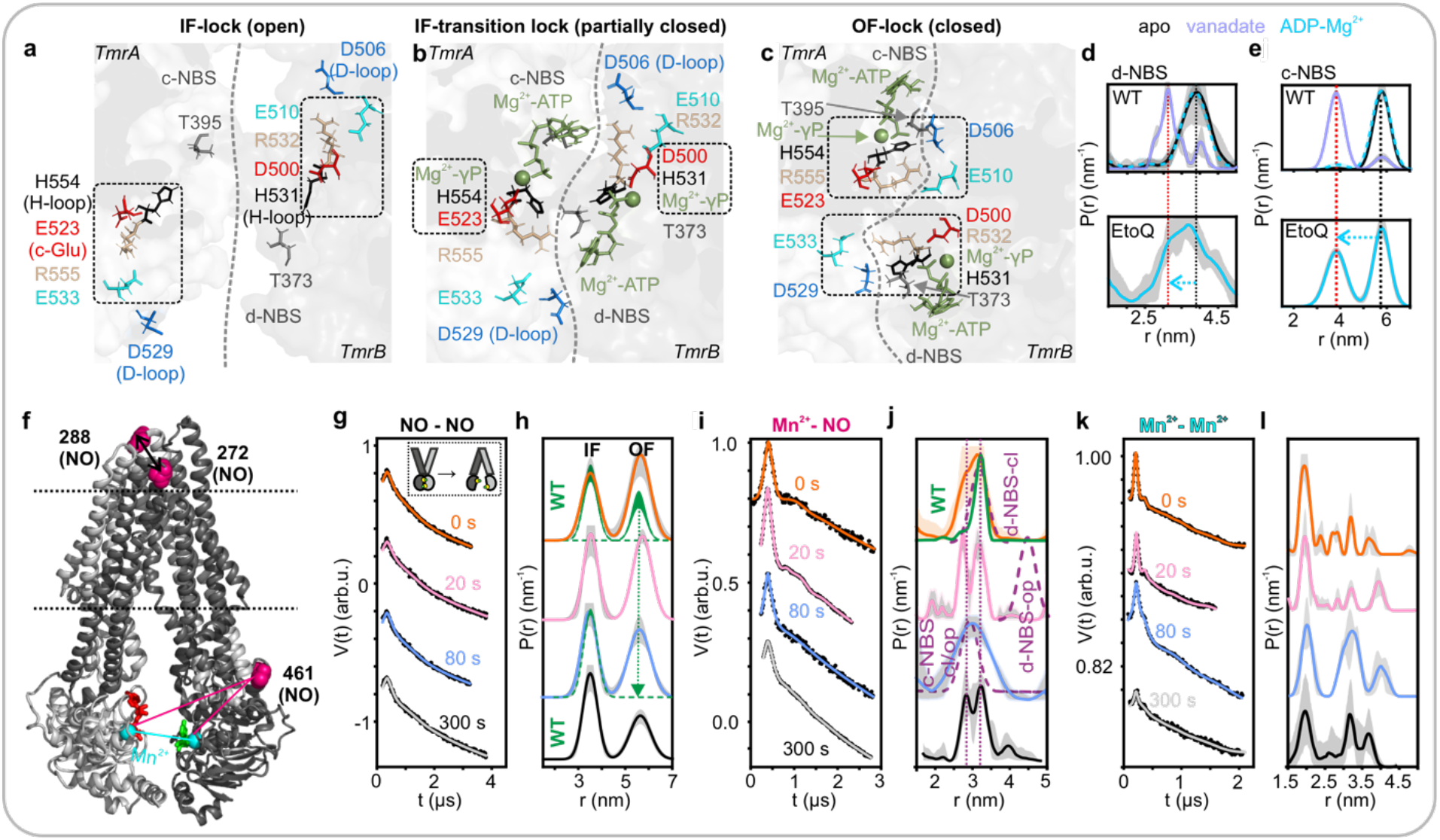
Energy coupling molecular locks and the reverse (OF → IF) transition in TmrAB. Snapshots from MD simulations of IF-apo, IF-Mg^2+^-ATP, and OF-Mg^2+^-ATP structures highlighting the residues forming the (**a**) IF-lock, (**b**) IF-transition lock, and (**c**) OF-lock in TmrA and TmrB (**Supplementary figures 19-21**). Each condition was sampled for 10 × 500 ns independent simulations. (**d– e**) Effect of Mg^2+^-ADP in the E→Q variant on the d-NBS and the c-NBS (**Supplementary figures 7g-h**). The apo and vanadate-trapped states (**Supplementary figures 2–3**) are overlaid as the reference for open and closed conformations, respectively. (**f-l**) Triple labeling and orthogonal PELDOR to observe the reverse transition in the E→Q variant. (**f**) The MTSL(NO)-labeled positions, bound nucleotides (in green and red), and the Mg^2+^ atoms (cyan) are highlighted in the IF structure (PDB ID: 6RAG). (**g–l**) NO - NO, Mn^2+^ - NO, and Mn^2+^ - Mn^2+^ PELDOR data and the corresponding distance distributions. The inset in panel (g) shows the observed reverse transition. The NO-NO data were globally analyzed based on an experimentally determined model (**Supplementary figure 1**). The NO-NO distances observed for the parental protein are overlaid in green (in panel h) (*14*). The Mn^2+^ - NO distance simulations in the open (op) and closed (cl) conformations are overlaid in panel (j). From position 461(NO), the Mn^2+^ - NO distance is unaltered at the c-NBS between the open and closed conformations (denoted as c-NBS cl/op in panel e).

The simulations further revealed that Mg^2+^-ATP binding (in the IF conformation) partially breaks the IF-lock and the NBSs move into a semi-closed conformation, forming an intermediate lock (named the IF-transition lock, **Figure 3b, Supplementary figures 21a–b)**. The TMDs remain closed during this transition. This intermediate structure is similar to the IF-narrow structure (PDB ID: 6RAF; RMSD = 2.46 Å), but having the NBDs more closed (*30*). At the c-NBS in TmrA, the c-Glu (TmrA-E523) and the H-loop TmrA-His554 moved together towards Mg^2+^−γ-phosphate, thereby detaching the c-Glu from TmrA-Arg555 and breaking the IF-lock (**Figure 3b**). An analogous interaction by TmrB-D500 (corresponding to c-Glu) and the H-loop TmrB-His531 with Mg^2+^−γ-phosphate is also formed at the d-NBS in TmrB.

Simulations with the Mg^2+^-ATP-bound OF structure showed that at the c-NBS in TmrA, the c-Glu interacts with Mg^2+^, and TmrA-His554 interacts with γ-phosphate and also forms an interfacial salt bridge with TmrB-Asp506 (D-loop Asp) from the opposing monomer (**Figure 3c)**. The neighboring TmrA-Arg555 residue also forms another interfacial salt bridge with TmrB-Glu510. The D-loop TmrB-Asp506 residue acts as a central hub by further interacting with TmrA-Thr395 in the Walker A motif (GXT_395_XXGKT]). The latter also interacts with the γ-phosphate, thereby indirectly connecting TmrB-Asp506 with the γ-phosphate as well. Thus, in the OF conformation, TmrA-Thr395, TmrA-Glu523, TmrA-H554, TmrA-R555, TmrB-D506, and TmrB-E510 together form a second molecular lock (named the OF-lock) at the c-NBS, with Mg^2+^− γ-phosphate as the pivot. The identical residues (but from opposite monomers) form the OF-lock at the d-NBS as well (**Figure 3c**). For the OF-lock, the interactions coordinated by Mg^2+^-γ-phosphate are primary; their disruption upon hydrolysis can act as the trigger for the reverse transition. At both NBSs, the core residues that form IF-lock or OF-lock also form an outer shell of weak non-ionic interactions, which might be important for overall stabilization (**Supplementary figures 19-20)**.

A conservation analysis with TmrA and TmrB as the query sequences revealed that all the residues involved in IF-lock and OF-lock are conserved across different classes of ABC proteins (**Supplementary figures 19–20, 22)**. The D-loop Asp (TmrA D529 and TmrB D506), H-loop His (TmrA H554 and TmrB H531), c-Glu (TmrA E523), and a hydrogen-bond-donating residue within the Walker A motif (TmrA T395 and TmrB T373) are conserved and essential among ABC transporters (*40, 41*). These residues are invariant among ABC exporters, including the eukaryotic transporters. On the other hand, the Arg next to the H-loop histidine (TmrA R555 and TmrB R532) and the Glu (or occasionally a negative residue) next to the D-loop (TmrA E533 and TmrB E510) are conserved only among type IV and type VI ABC transporters and the distantly related DNA repair soluble ABC protein Rad50, suggesting that they play a less critical role compared to other residues.

As c-Glu is the key residue for the IF-lock at the c-NBS (**Figure 3a**), the E→Q substitution can weaken the IF-lock and favor IF → OF transition. This might explain why in the E → Q variant, ATP alone (without Mg^2+^) can increase the initial burst and the *K*_*eq*_ for TMDs opening (**Figure 2d)**. Remarkably, ADP-Mg^2+^ alone can close the NBSs in a fraction of the transporters in the E → Q variant (**Figure 3d–e, Supplementary figures 7g–h**), substantiating that destabilizing the IF-lock is one of the deterministic steps for the IF → OF transition. As the ATPase activity is more than 1000-fold reduced in the E→Q variant (*42*), it is an ideal background for elucidating the role of ATP hydrolysis for the reverse (OF → IF) transition. We observed the OF → IF transition by concomitantly monitoring the TMDs and the NBSs within the same molecules (**Figures 3f–l**). For this purpose, we used Mn^2+^, which can functionally replace Mg^2+^ (*14*). We attached two MTSL labels at the TMDs and a third one near the d-NBS (at TmrA 461, **Figure 3f**) and determined NO - NO (TMDs), Mn^2+^ - NO (NBSs), and Mn^2+^ - Mn^2+^ (Mn^2+^ -ATPs) distances in a time-resolved manner while the transporter moves from OF to the IF conformation (see methods). During this reverse transition, TMDs close, associated with the opening of the NBSs. The NO - NO data revealed a gradual reversal from OF to the IF conformation (**Figures 3g–h**). However, a considerable fraction of the TMDs remained in the OF conformation, whereas the parental protein fully reversed in an accelerated manner (*14*) (**Figure 3h**, green dotted lines).

The Mn^2+^ - NO experiments revealed a broadened distribution corresponding to a ∼1:1 closed conformation of both NBSs (**Figures 3i–j)**. As with the TMDs, the sample also contained an open population for both NBSs, which is spectroscopically invisible (see methods). With the parental protein (**Figure 3j**, green line), distances corresponding to the c-NBS were already absent at the very beginning due to ATP hydrolysis (*14*). The modulation depth gradually decreased towards 300 s, indicating a slow release of Mn^2+^-ATP and the opening of both NBSs. The Mn^2+^ - Mn^2+^ PELDOR (**Figures 3k–l**) revealed a major peak around 2.0 nm in line with the Mg^2+^ - ATP-bound OF structure (PDB ID: 6RAH). These measurements are not possible in the parental protein due to fast ATP hydrolysis and Mn^2+^ release at the c-NBS (*14*). We observed additional, longer distances that became pronounced at 80 s, suggesting a novel intermediate state during dissociation of Mn^2+^ - ATP. The modulation depth was further decreased towards 300 s, revealing stepwise Mn^2+^-ATP dissociation from the NBSs (see methods). In summary, the data reveal that in the E → Q variant, the forward transition is enhanced (**Figure 2d**), whereas the reverse transition is significantly slower (**Figures 3h, j, l**).

We concluded that the reverse transition is slower in the E→Q variant because ATP hydrolysis is necessary for this step. The P_i_ might be immediately released through exit tunnels following hydrolysis, before ADP dissociates and the NBSs open (*14, 30, 43*). As γ-phosphate acts as the pivot (**Figure 3c**), its release might destabilize the OF-lock. We used γ-phosphate/P_i_ removal from Mg^2+^ - ATP at the c-NBS as a nonequilibrium perturbation to destabilize the OF state and probe the early responses associated with reverse transition using MD simulation. Strikingly, this induced a rapid opening of the c-NBS (**Figure 4a, Supplementary figure 24**), in agreement with a strained conformation of the c-NBS as suggested by MFPT calculations (**Figures 2e–g**). These (and the other changes as detailed below) were not observed in the parental TmrAB when the γ-phosphate was not removed or in the E→Q variant, even after the γ-phosphate removal (**Figures 4a–c, d–g**). Notably, upon γ-phosphate removal, all the key interfacial interactions that stabilize the OF-lock are lost (**Figure 4d–g, Supplementary figure 23**), and the transporter gradually reforms the IF-lock at the c-NBS, accompanied by its opening (**Figure 4i–j, Supplementary figures 23i–m**). Also, the Mg^2+^ ion moves between the α- and β-phosphates of ADP at the c-NBS (**Supplementary figure 23l**), abolishing its interaction with c-Glu (TmrA-E523) and destabilizing the OF-lock. These changes are accompanied by an increased flexibility of the TMDs at the periplasmic side, approaching the occluded conformation during the simulation time (**Figure 4h, Supplementary figure 24c**).

**Figure 4.**
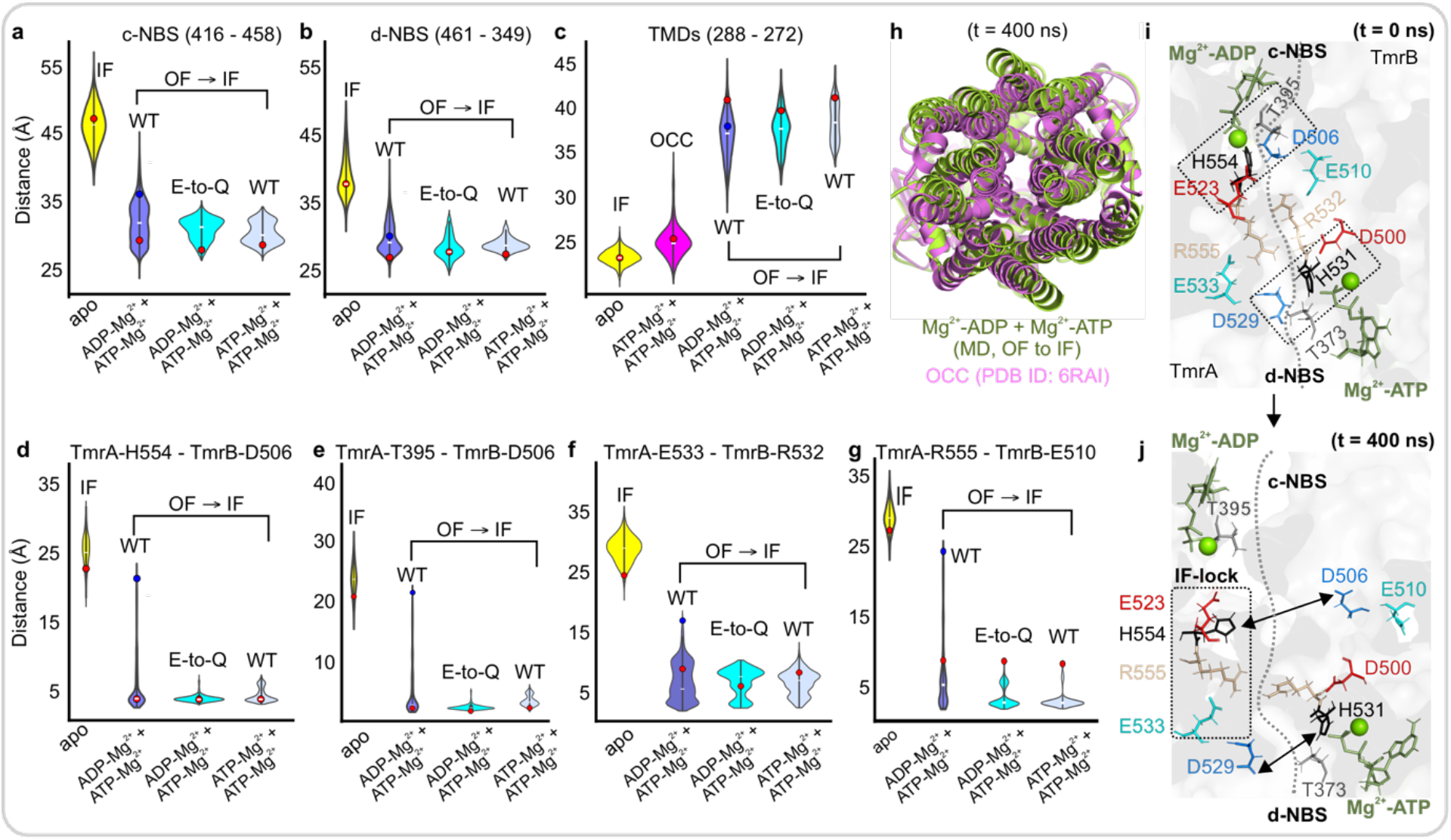
Molecular dynamics simulation of reverse transition (OF → IF) in the post-hydrolytic state. The γ-phosphate was manually removed from the c-NBS before the simulation, and the E→Q variant was used as the direct control, along with WT TmrAB without γ-phosphate removal. (**a-c**) Violin plots showing probability distribution of the c-NBS (**a**), d-NBS (**b**), TMDs (**c**), and the interfacial interactions at the c-NBS (**d-g**) as indicated. Minimum heavy-atom distance distributions for the residue pairs were sampled from 10 × 500 ns MD simulations (see **Supplementary figures 23–24**). (**h**) A top-view of the overlay of the TMDs for the OCC structure and the intermediate state from MD during the OF → IF transition and the corresponding rearrangement of the OF-locks (**i–j**) at the two NBSs.

Eventually, the d-NBS also opens (**Figure 4b, j, Supplementary figures 23m, 24a**), making it one of the slower steps during reverse transition. Thus, while c-NBS closure acts as the primary kinetic barrier during the forward transition, ATP hydrolysis shifts the bottleneck to the d-NBS. The two NBSs and the periplasmic side of the TMDs independently fluctuated during simulation, but the TMDs did not fully restore to the IF conformation; in particular, the cytosolic gate remained stable (**Supplementary figures 24c–d)**. Thus, opening of the cytosolic gate at the TMDs might constitute the key barrier for the completion of the reverse transition to the IF conformation. Absence of all of these changes in the E→Q variant (in agreement with the delayed kinetics revealed from PELDOR data, **Figure 3f-I**) shows that P_i_ release alone is insufficient, and the restoration of the IF-lock at the c-NBS (**Figure 4j**, for which c-Glu is the key residue) is also necessary to trigger c-NBS opening and reverse transition.

## Conclusions

Structural, biochemical, and biophysical studies have provided a detailed understanding of the conformational cycle driving substrate transport in ABC transporters (*31, 44-46*). Our integrated framework resolves the mechanism of this chemomechanical transduction, identifying a pair of conserved ionic locks within the nucleotide-binding sites (NBSs) that function fundamentally as dual-action energy coupling modules and discrete transition barriers. While the forward transition relies on ATP binding (but not ADP) to mechanically rupture the IF-lock, subsequent hydrolysis provides the driving force to overcome the OF-lock barrier and reset the cycle. Mg^2+^ acts as a key player within this landscape and accelerates the transition rates to physiologically relevant time scales. Crucially, this gating manifests as an asymmetric kinetic divergence between the c-NBS and TMDs, which undergo anti-correlated kinetic shifts in opposite directions during the forward and reverse stages of the transport cycle. The delayed c-NBS closure, as the limiting step for the forward transition, might ensure a sufficient window for substrate diffusion into the binding pocket (**Figure 5a–b**). Because the periplasmic gate closes earlier during the reverse transition, shifting the rate-limiting bottleneck to cytosolic TMD opening prevents substrate back-transport. It ensures strictly unidirectional translocation (**Figure 5c–e**), thereby breaking microscopic reversibility via kinetic control. Given the evolutionary conservation of these electrostatic coupling hubs, this interplay between ATP binding/hydrolysis, barrier gating, and mechanical transduction might represent a unifying physical principle underlying chemomechanical coupling across the broader superfamily of ABC transporters and related ATPases.

**Figure 5.**
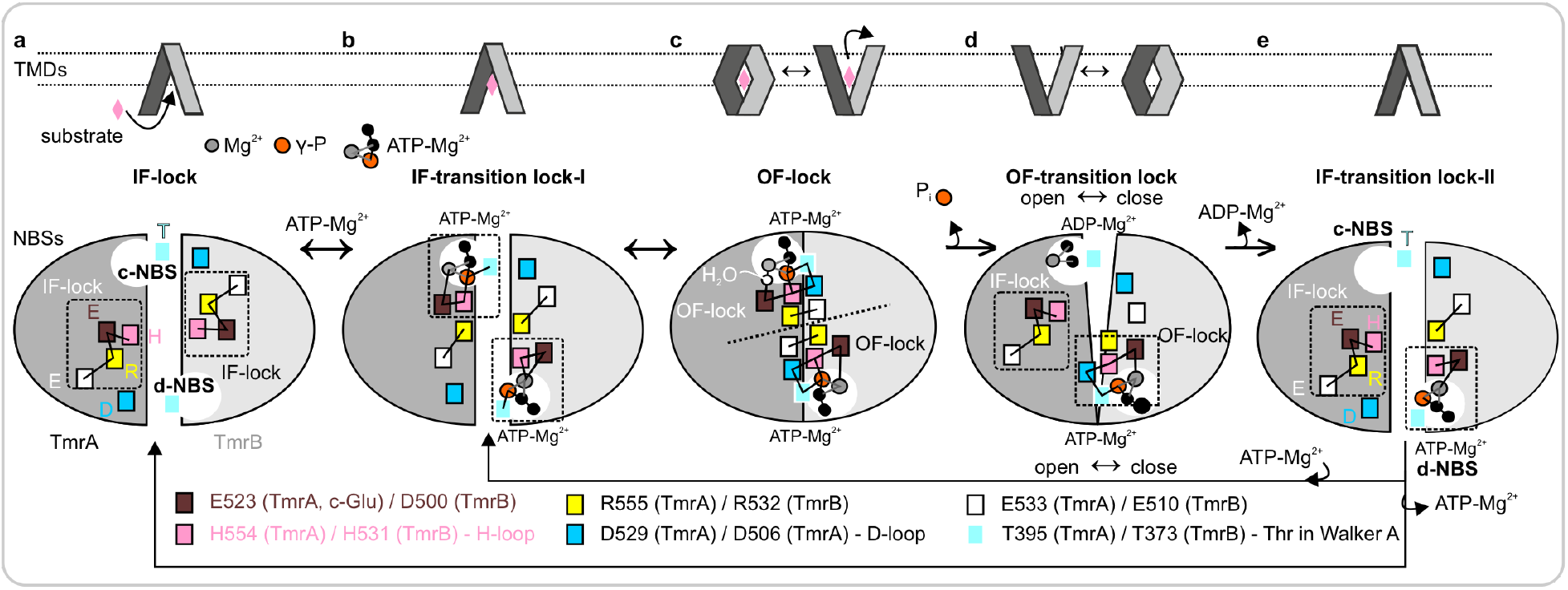
ATP-mediated chemomechanical energy transduction in TmrAB. (**a**) The IF conformation involves the IF-lock in both subunits, formed by four conserved residues (inside the box). The TMDs have an IF conformation in which the substrate binds inside the cavity. (**b**) ATP binding at the NBSs partially breaks the IF-locks, primarily by displacing the c-Glu and H-loop His toward Mg^2+^-γ-phosphate, forming the IF-transition lock-1 at the NBSs. (**c**) The OF-lock stabilizes the closed conformation with Mg^2+^- γ-phosphate as the pivot, TMDs are switched to OF conformation, and the substrate diffuses out. MD simulation shows that at the c-NBS, the c-Glu bridges a water molecule with the Mg^2+^ ion. (**d**) ATP hydrolysis releases the γ-phosphate (P_i_), thereby destabilizing the OF-lock and reforming the IF-lock at the c-NBS, leading to its opening and transition into an asymmetric OF-transition lock. This triggers the transition of the TMDs towards the OCC conformation and increases the dynamics of the d-NBS. The TMDs and the NBS exhibit independent flexibility. (**e**) Mg^2+^-ADP dissociates from the c-NBS, the d-NBS, and eventually the cytosolic gate at the TMDs open to complete the reverse transition. Depending on the nucleotide concentration, Mg^2+^-ATP may either dissociate from the d-NBS (**e–a**) or bind to the c-NBS (**e– b**) to restart the transport cycle.

## Acknowledgements

B. J. would like to thank Thomas F. Prsiner for providing the spectrometer time for a few of the samples.

## Funding

Emmy Noether program from German Research Foundation (DFG) JO 1428/1-1 (BJ)

Rise up! Grant from the Boehringer Ingelheim Foundation (BJ)

Collaborative Research Center (CRC1507/P18 and CRC1507/P02) funded by the German Research Foundation (DFG, B.J and R.T)

## Author contributions

Conceptualization: RN, BJ; Methodology: MR, HB, RT, RN, BJ: Investigation: MR, HB, MP, KB, CH; Visualization: MR, HB, BJ; Funding acquisition: BJ; Project administration: BJ; Supervision: RT, RN, BJ; Writing – original draft: MR, HB, BJ; Writing – review & editing: MR, HB, CH, RT, RN, BJ

### Competing interests

Authors declare no competing interests

### Data, code, and materials availability

All the primary experimental data corresponding to the PELDOR experiments are presented in the main and/or supplementary figures. The relevant MD trajectories are deposited in Zenodo (https://doi.org/10.5281/zenodo.20678086), which also contains the umbrella-sampling pullx files for all three collective variables and the MD simulation setup table (Supplementary Table 3). Additional data and analysis that support the conclusions of this work are available upon request to the authors.

## Notes

https://doi.org/10.5281/zenodo.20678086

### Competing Interest Statement

The authors have declared no competing interest.

