## Supplementary Materials for "Kinetic asymmetry drives directionality in an ATP-binding cassette transporter"

### Experimental Procedures

#### Cloning, expression, purification, and conservation analysis of TmrAB

TmrAB cysteine variants were prepared as previously described (*1, 2*). Protein was expressed in *E. coli* BL21(DE3) in LB medium at 37 °C. At an $OD_{600}$ between 0.6 and 0.8, expression was induced by the addition of 0.5 mM isopropyl β-D-thiogalactopyranoside (IPTG, Roth). Cells were grown for 4 h at 37 °C. Harvested cells were resuspended in lysis buffer A (20 mM HEPES pH 7.5, 300 mM NaCl, 50 µg $ml^{-1}$ lysozyme), disrupted by sonication (Branson sonifier 250, Thermo Scientific) and cell debris was removed by centrifugation at 13500x*g* for 15 min at 4 °C. Membranes were pelleted at 200000x*g* for 1 h at 4 °C and solubilized in purification buffer (20 mM HEPES pH 7.5, 300 mM NaCl, 20 mM n-dodecyl-β-D-maltoside (β-DDM)) for 1 h at 4°C. Insoluble components were removed by centrifugation at 200000x*g* for 30 min at 4 °C. Solubilized TmrAB was incubated with Ni-NTA agarose (Qiagen) with 1 mM β-mercaptoethanol (β-ME) and 30 mM imidazole for 1 h at 4 °C. After washing with 10 column volumes of washing buffer (20 mM HEPES pH 7.5, 300 mM NaCl, 50 mM imidazole, 1 mM β-ME, 1 mM β-DDM), bound TmrAB was eluted with elution buffer (20 mM HEPES pH 7.5, 300 mM NaCl, 300 mM imidazole, 1 mM β-DDM) and buffer exchanged in a PD-10 desalting column (Cytivia) to 20 mM HEPES pH 7.5, 150 mM NaCl, 1 mM β-DDM. For conservation analysis of the key residues directly involved in the molecular locks (Figure 3a-c), related sequences were fetched using TmrA and TmrB as the query sequences against the Uniprot database using the PSIBLAST tool[1]. Multiple sequence alignment was performed with the first 500 sequences showing the closest match using Clustal Omega (*3*). In a second step, an additional analysis was performed for the partner residues that interact with the above key residues. Here, homologous sequences of TmrA and TmrB were retrieved independently from UniProt using a 50% sequence identity threshold. This yielded 91 sequences for TmrA and 48 sequences for TmrB. The retrieved sequences were aligned separately for each subunit and used for subsequent conservation analysis.

#### Reconstitution into nanodiscs, liposomes, and transport activity

For nanodisc reconstitution, labeled TmrAB 288-272, membrane scaffold protein MSP1D1, and bovine brain lipid extract were combined in a molar ratio of 1:7.5:200. The solution was then diluted with TSK buffer and incubated for 30 min at RT on the roller mixer. After 30 min, 200 mg bio-beads in TSK buffer per 500 µl sample were added. Following incubation at 4 °C for 1 h, the bio-beads were exchanged, and the mixture was incubated at 4 °C overnight. The bio-beads were removed afterwards, and the solution was centrifuged for 3 min at 3000x*g* and subsequently loaded onto a Superdex 200 Increase10/300 GL column (GE Healthcare). The fractions corresponding to nanodisc-reconstituted TmrAB were pooled and concentrated to 15 – 30 µM. For liposome reconstitution, *E. coli* polar lipids/DOPC (7:3) (Avanti Polar Lipids) with a total lipid concentration of 20 mg $mL^{-1}$ were prepared in the buffer (20 mM HEPES, pH 7.5, 140 mM NaCl, 5% (v/v) glycerol). The liposomes were extruded 11 times through a 400 nm polycarbonate filter. After treating liposomes for 30 min at 4°C with Triton X-100 (Roth), spin-labeled TmrAB was added at a molar protein-to-lipid ratio of 1:15,000 for transport assays. The sample volume was adjusted to a final lipid concentration of 10 mg $mL^{-1}$. Following incubation for 30 min at 4°C, the detergent was removed stepwise using Bio-Beads (Bio-Rad). Proteoliposomes were pelleted by ultracentrifugation at 280,000x*g* for 30 min at 4°C. Pelleted proteoliposomes were resuspended in reconstitution buffer to a final lipid concentration of 5 mg $mL^{-1}$, and reconstitution efficiency ranged between 40 and 50%. For transport assays, proteoliposomes (1 mg $mL^{-1}$ lipid final) were incubated with 3 mM ATP, 3 mM $MgCl_2$, and 3 µM fluorescent peptide substrate (RRYCFLKSTEL) (*4*) in transport buffer (20 mM HEPES pH 7.5, 100 mM NaCl, 5% (v/v) glycerol) for 5 min at 68 °C. Reactions were stopped with four volumes of ice-cold stop buffer (10 mM $Na_2HPO_4$, 1.8 mM $KH_2PO_4$, pH 7.3, 140 mM NaCl, 2.7 mM KCl, and 10 mM EDTA). Samples were transferred to a Multi Screen Filter Plate (Durapore membrane 0.65 µm; Millipore) and washed twice with five volumes of stop buffer. The liposomes were lysed using 1 × PBS supplemented with 0.1% of SDS for 10 min. The samples were transferred into a microtiter plate, and fluorescence was quantified using a microplate reader ($\lambda_{ex}/\lambda_{em}$ 485/520 nm).

#### Spin-Labeling of TmrAB and CW EPR Spectroscopy

Cysteine variants of TmrAB were labelled with a 40-fold molar excess of methanethiosulfonate spin label (1-oxyl-2,2,5,5-tetramethylpyrroline-3-methyl, MTSSL) in sample buffer for 30 min at room temperature. The protein was concentrated using Vivaspin 6 filters with a 50 kDa cutoff (Merck), and free MTSSL was removed by Micro Bio-Spin (Biorad). Labelling efficiency was determined by continuous wave (cw) EPR measurements on a Bruker EMXNano spectrometer with the following experimental parameters: 100 kHz modulation frequency, 1.5 G modulation amplitude, 1 mW microwave power, 5.12 ms time constant, 22.5 ms conversion time, and 15 mT sweep width.

#### Sample Preparation for PELDOR / DEER spectroscopy

All samples were prepared in TSK buffer (20 mM HEPES pH 7.5, 150 mM NaCl, 1 mM β-DDM). For determination of the distances in the reference states at the NBSs and the TMDs (open or closed conformation), 50 µM TmrAB was prepared either under apo-condition (without any nucleotide), or in the $Mg^{2+}$-ADP-$VO_4^{3-}$ -trapped state (10 mM ATP, 10 mM $Na_3VO_4$, 10 mM

$MgCl_2$, 15 % $d_8$ glycerol, 50 °C, 5 min), and frozen in liquid nitrogen. For the time-resolved PELDOR experiments (IF ↔ OF), samples were prepared either under ATP-EDTA conditions (50 μM TmrAB, 50 mM ATP, 0.5 mM EDTA, in Figure 1e-g) or turnover conditions (50 μM TmrAB, 50 mM ATP, 50 mM $MgCl_2$, in Figure 2b-c), incubated at different temperatures (30 - 65 °C) in preheated sample tubes for incremental times (between 10 - 600 s) and immediately frozen in liquid nitrogen. The sample temperatures within the tested range were reached within 2–3 s, and subsequent freezing took 1 - 2 s under the experimental setup. For the time-resolved observation of reverse transition (Figure 3f-l), samples were prepared in the presence of ATP-$Mn^{2+}$ (65 μM TmrAB, 10 mM ATP, 10 mM $MnCl_2$, 15 % glycerol), incubated at 50 °C for 5 min, and subsequently transferred to ice. Free $Mn^{2+}$ and ATP were removed using Micro Bio-Spin columns (Bio-Rad) equilibrated with ice-cold TSK buffer and 0.5 mM EDTA. Afterwards, 15 % deuterated glycerol was added and frozen immediately (time point 0 s) or transferred to preheated quartz tubes, incubated at 50 °C for increasing times (up to 300 s), and frozen immediately in liquid nitrogen. The first sample (time point 0 s) contains an equilibrium between open and closed conformations of the NBSs and the TMDs (a fraction of the transporters reverses during the Bio-Spin step following incubation with ATP-$Mn^{2+}$). This is evident for the TMDs (Figure 3g-h). For the NBSs, the $Mn^{2+}$-NO experiments (Figure 3i-j) cannot detect the open NBS, as $Mn^{2+}$-ATP dissociates in this conformation, especially since the sample has no free ATP after desalting. For $Mn^{2+}$-$Mn^{2+}$ experiments (Figure 3k-l), the 80 s sample showed an increased fraction of a novel longer distance component, representing an intermediate state (between $Mn^{2+}$-$Mn^{2+}$ centers). This manifested as an apparent increase in modulation depth as the shorter distances (centered at 1.9 nm) in other samples were not fully excited (and thereby reduced the effective modulation depth). These longer $Mn^{2+}$-$Mn^{2+}$ distances were not present in the corresponding $Mn^{2+}$-NO experiments (Figure 3i-j, 80 s), implying that those must correspond to $Mn^{2+}$ binding mode too far to be observed from position 461 at which nitroxide label is attached (Figure 3f).

#### PELDOR / DEER Spectroscopy, data analysis, and simulations

PELDOR/DEER measurements were performed on a Bruker Elexsys E580 Q-Band Pulsed spectrometer equipped with an arbitrary waveform generator (SpinJet-AWG), a continuous-flow helium cryostat, a temperature control system (Oxford Instruments), a 50 W solid state amplifier, and a Bruker EN5107D2 dielectric resonator. Echo-detected ESR spectrum and $T_M$ measurements were performed at 5 K for $Mn^{2+}$ or at 50 K for nitroxide using the two-pulse Hahn echo sequence $\pi/2 - \tau - \pi - \tau$ employing Gaussian pulses (48 ns) and an inter-pulse delay ($\tau$) of 200 ns. The phase memory time ($T_M$) values were determined by systematically varying the $\tau$ and fitting of the data to a stretched exponential function ($\exp[-(2\tau/T_M)^{\kappa}]$). The $T_M$ was invariant between the open/closed conformations at the TMDs and NBDs under the experimental conditions (in Figure 1e-g and Figure 2b-d), such that no relaxation-dependent correction was necessary for the determined probabilities of the populations. The four-pulse PELDOR experiments were recorded using Gaussian pulses and a dead-time free sequence employing a 16-step phase cycling (x[x][xp]x) (*5*). Nitroxide – nitroxide measurements were acquired at 50 K using a 38 ns pump pulse and 48 ns observer pulses. The pump pulse was set to the maximum of the echo-detected field-swept spectrum, while the observer pulses were set at 80 MHz lower, and the shot repetition time (SRT) was kept at 2 ms. Nitroxide–$Mn^{2+}$ PELDOR experiments were recorded at 5 K while pumping the nitroxide with a 16 ns pulse and observing the $Mn^{2+}$ with 24 ns Gaussian observer pulses. The observer pulses were set at -248.4 MHz away from the pump pulse. The SRT was kept at 1 ms. $Mn^{2+}$–$Mn^{2+}$ PELDOR experiments were recorded at 5 K. The pump pulse was set at +259 MHz away from the observer pulses. A 12 ns pump pulse and 24 ns observer pulses (both Gaussian) were employed. The nitroxide – nitroxide five-pulse PELDOR/DEER experiments were performed according to the pulse sequence $\pi/2_{obs} - (\tau/2 - t_0) - \pi_{pump} - t_0 - \pi_{obs} - t' - \pi_{pump} - (\tau - t + \delta) - \pi_{obs} - (\tau_2 + \delta)$ with the $\delta$ set to 200 ns. Experiments were performed at 50 K using 48 ns Gaussian observer pulses and 16-step phase cycling ($xx_p[x][x_p]x$) (*5*). A 36 ns standing and a 48 ns moving pump pulse (both Gaussian) were employed. Distance distributions were determined using the DeerLab program[5]. The primary data were simultaneously fitted for distances and the background function using a model-free Tikhonov regularization (TR) or a Gaussian model-based approach, as indicated. For related data sets (as in Figures 1e-g and Figure 2b-d), global analysis was performed. Simulations of the distance distributions were obtained on the corresponding structures using the rotamer libraries for MTSL as implemented in the MATLAB-based software package MMM2022.2[6].

#### Kinetic and thermodynamic analysis

All the analyses were performed using Python 3.0 packages. The temporal evolution of the populations, as determined from TR-PELDOR, was further fitted to a first-order kinetic relaxation according to $P(t) = P_{eq} - (P_{eq} - P_0) * e^{-k_{tot}*t}$. The forward ($k_1$) and reverse ($k_{-1}$) rate constants were calculated as $P_{eq}*k_{tot}$ and $(1-P_{eq})*k_{tot}$, respectively. The uncertainties were calculated using first-order Gaussian error propagation, including covariance terms from the nonlinear fit. The determined rate constants were fitted to Kramers' reaction rate theory as $ln(k) = [(-\Delta H^{\ddagger}_{structural} + E_{\eta}) / R]\ (1/T) + ln(A_{molecular} / \eta_0)$. $\Delta H^{\ddagger}_{structural}$ is the intrinsic structural activation enthalpy barrier of the protein domain transition, $E_{\eta}$ is the activation energy of viscous fluid flow for the solvent/micelle environment, fixed as a known physical constant of 17 kJ·mol$^{-1}$, $A_{molecular}$ is the effective kinetic pre-exponential factor (frequency factor), and $\eta_0$ is the temperature-independent baseline viscosity of the solvent medium. Experimental uncertainties of the raw rate constants were propagated into logarithmic space using a first-order Taylor series expansion, where the standard deviation for each data point was defined as $\sigma_{lnk}$ = $SE_k$ / $k$. These transformed errors were utilized as individual weighting factors in a weighted linear least-squares regression analysis to determine the optimized slopes and intercepts. The net transition enthalpy ($\Delta H^0$) and entropy ($\Delta S^0$) governing the ground-state equilibrium were calculated directly from the differences between the forward and reverse kinetic pathways, which effectively cancels out isotropic environmental solvent-drag parameters ($E_\eta$ and $\eta_0$). Uncertainties for these net thermodynamic endpoints were subsequently determined through standard Gaussian error propagation using the independent standard errors extracted from the variance-covariance matrices of the univariate regressions for each kinetic direction.

#### Molecular dynamics simulations

Cryo-EM structures of the heterodimeric ABC exporter TmrAB in three conformational states were used as the starting model(s) as indicated: inward-facing wide (PDB ID: 6RAG), ATP-bound outward-facing open (PDB ID: 6RAH), and occluded (PDB ID: 6RAI). Missing side chains were rebuilt using CHARMM-GUI. Nucleotides were retained as resolved in the experimental structures unless stated otherwise. For the inward-facing model, we aimed to obtain ATP bound at both nucleotide-binding sites; therefore, the ADP resolved at the c-NBS in 6RAF (which contains ATP-$Mg^{2+}$ at the d-NBS) was replaced with ATP-$Mg^{2+}$ using the relevant coordinates from the outward-facing structure 6RAH. For the outward-facing model, to mimic the post-hydrolytic state, the γ-phosphate was removed from the ATP-$Mg^{2+}$ molecule at the c-NBS to produce an ATP/ADP-bound state. The coordinates of the modified nucleotide-binding site were subsequently refined by energy minimization, which resulted in a shift of the $Mg^{2+}$ ion toward the α- and β-phosphates. The systems were solvated with TIP3P water molecules and neutralized with NaCl at a final ionic strength of 0.15 M. All simulations were performed with GROMACS 2024, using the CHARMM36m force field (*6, 7*). Energy minimization was followed by equilibration under NVT and NPT ensembles, gradually releasing position restraints on protein and lipid heavy atoms. Long-range electrostatics were calculated with the particle mesh Ewald method (*8*) using a real-space cutoff of 12 Å. van der Waals interactions were smoothly shifted to zero between 10 and 12 Å. All covalent bonds involving hydrogen were constrained with the LINCS algorithm (*9*), allowing a 2 fs integration step. The velocity-rescaling thermostat (*10*) was applied to maintain temperature, and the Parrinello–Rahman barostat (*10*) controlled pressure. Umbrella sampling simulations were performed up to 500 ns per window at 310 K. Production simulations were carried out at 333 K and 1 bar. IF-start simulations were run for 10 x 500 ns per system, and OCC-start simulations were run for 6 x 1.7 μs per system. Umbrella-sampling windows were simulated for 500 ns per window at 310 K.

#### Umbrella Sampling and Potential of Mean Force (PMF)

To characterize the free-energy landscape along the chosen reaction coordinate (x), we performed umbrella sampling simulations using GROMACS 2025.2. A harmonic biasing potential with a force constant of 1000 kJ $mol^{-1}$ $nm^{-2}$ was applied to restrain the collective variable at different target values, producing an ensemble of overlapping windows. Each trajectory was saved as a pullx time series containing the time evolution of the restrained coordinate. The unbiased potential of mean force (PMF) / *W(x)*, was reconstructed from these data with the weighted histogram analysis method (WHAM).

#### Mean First-Passage Time Calculation

To estimate the transition timescale along the reaction coordinate, we employed the one-dimensional Fokker-Planck equation

$$\frac{\partial P(x,t)}{\partial t} = \frac{\partial}{\partial x}\left\{\left[\frac{k_B T}{\gamma_{eff}(x)}\right] e^{-U(x)/(k_B T)} \frac{\partial}{\partial x}\left[P(x,t) e^{U(x)/(k_B T)}\right]\right\} \quad (1)$$

where P(x,t) is the probability distribution of the reaction coordinate x, $U_{(x)}$ is the potential of mean force (PMF), $k_{BT}$ is the thermal energy at 310 K, and $\gamma_{eff}^{(x)}$ is the position-dependent effective friction coefficient. For a process originating at $x_s$ and terminating at an absorbing boundary $x_f$, the mean first-passage time (MFPT) reads

$$\tau^{MFP}(x_s, x_f) = \int_{x_s}^{x_f} dx \left[\frac{\gamma_{eff}(x)}{k_B T}\right] e^{U(x)/(k_B T)} \int_{-\infty}^{x} dx'\, e^{-U(x')/(k_B T)} \quad (2)$$

for $x_s < x_f$. A reflecting boundary condition is imposed at $x_{left}$, placed at the edge of the sampled PMF region, which sets the lower limit of the inner integral. The PMF U(x) was obtained from umbrella sampling simulations combined with the multistate Bennett acceptance ratio (MBAR) estimator. The MFPT integral was evaluated numerically using the trapezoidal rule over the full PMF grid.

Memory effects are universally present in biomolecular dynamics and influence barrier-crossing timescales. To account for these effects, the position-dependent effective friction $\gamma_{eff}^{(x)}$ was extracted directly from the umbrella sampling trajectories using a recently derived approach (*11, 12*) based on the position autocorrelation function. For each umbrella window centred at the mean position ⟨x⟩, the position autocorrelation function is defined as

$$C^{xx}(t) = \langle (x(0) - \langle x \rangle)(x(t) - \langle x \rangle) \rangle \quad (3)$$

The effective friction at the mean window position is then given by

$$\gamma_{eff}(x = \langle x \rangle) = \frac{2k_BT}{[C^{xx}(0)]^3} \int_0^\infty dt [C^{xx}(t)]^2 \quad (4)$$

This expression effectively incorporates non-Markovian memory effects into the barrier-crossing time through the full time integral of the squared autocorrelation function. To evaluate the integral in Eq. 4 analytically and avoid numerical integration of noisy long-lag data, the autocorrelation function of each window was fitted to a sum of N exponential decays,

$$C_{fit}(t) = \sum_i \frac{g_i}{\tau_i} e^{-t/\tau_i} \quad (5)$$

where gi and τi are the amplitude and decay time of the i-th exponential component, respectively. The fit was performed over a log-spaced subsample of the autocorrelation in the range where signal exceeds noise. Substituting Eq. 5 into Eq. 4 yields the analytical expression

$$\int_0^\infty dt \left[C_{fit}(t)\right]^2 = \sum_i \frac{{g_i}^2}{2\tau_i} + 2 \sum_{i<j} \frac{g_i g_j}{\tau_i + \tau_j} \quad (6)$$

The convergence of this integral was verified for each umbrella window by computing the running integral I(t)/I(∞) and confirming that it reaches a plateau well before the autocorrelation truncation time. Windows in the barrier region carry the dominant weight in the MFPT integral through the Boltzmann factor $\exp[U_{(x)}/k_{BT}]$ and showed integral convergence values of 0.94–1.00 at the truncation time (Supplementary figure 14). The running integral of $[Cxx(t)]^2$ that enters Eq. 4 is shown for every umbrella window in Supplementary figure 13. The discrete effective friction values $\gamma_{eff}^{(x)}$ obtained per umbrella window were used to construct a continuous friction profile by fitting to the sigmoid function

$$log_{10}\gamma_{eff}^{fit}(x) = \frac{\sigma}{\left[1+e^{(x-x_0)/w}\right]} + c \quad (7)$$

where σ is the amplitude of the sigmoid drop, $x_0$ is the midpoint of the friction transition, w is the width parameter, and c is the asymptotic baseline value of $\log_{10}(\gamma)$ at large x. The fitted continuous profile $\gamma^{fit}{}_{eff}{}^{(x)}$ was used as the primary input to Eq. 2.

For comparison, two further friction estimators were computed from the same umbrella-sampling windows (Supplementary figure 17a). The position-autocorrelation friction $\gamma_{pos}(x)$ is the Markovian, single-relaxation estimate of Hummer(*13*),

$$\gamma_{pos} = (K^2/k_BT) \int_0^\infty C^{xx}(t)\, dt \quad (8)$$

where $K = k_BT/\langle(x-\langle x\rangle)^2\rangle$ is the variance-based stiffness of the effective harmonic well in each window. With the multi-exponential fit of $C^{xx}(t)$ (Eq. 5) the integral evaluates analytically to the sum of the fit amplitudes $\Sigma_i\, g_i$ (Supplementary figure 12). Because $\gamma_{pos}$ weights the linear autocorrelation integral whereas $\gamma_{eff}$ (Eq. 4) weights the squared one, the two coincide for a single-exponential $C^{xx}(t)$ and differ only when $C^{xx}(t)$ is genuinely multi-exponential (Supplementary figure 11); the difference between $\gamma_{eff}$ and $\gamma_{pos}$ therefore quantifies the contribution of memory.

The solvent-force friction $\gamma_{sol}(x)$ follows the Green–Kubo route of Daldrop, Kowalik, and Netz (*14*),

$$\gamma_{sol} = (1/k_BT) \int_0^\infty \langle\delta F(0)\delta F(t)\rangle\, dt \quad (9)$$

where the instantaneous force on the restrained coordinate is $F(t) = m_{eff}\, a(t) + K(x(t)-\langle x\rangle)$, the effective mass is fixed by equipartition, $m_{eff} = k_BT/\langle v^2\rangle$, and v(t) and a(t) are obtained by Savitzky–Golay differentiation of the trajectory. The integral was evaluated from a multi-exponential fit of $\langle F(0)F(t)\rangle$ ($n_{exp}$ = 3, Supplementary figures 15–16). This estimate reflects the instantaneous (high-frequency) friction and, together with $\gamma_{pos}$, brackets the memory-corrected $\gamma_{eff}$ (Supplementary figure 17a). The effective friction $\gamma_{eff}$ (Eq. 4) is smaller, since it includes the effect of memory on the barrier crossing time, which for intermediate memory time has been shown to decrease the barrier-crossing time (*15, 16*). The first passage times following from the different friction estimates are shown in Supplementary figure 18.

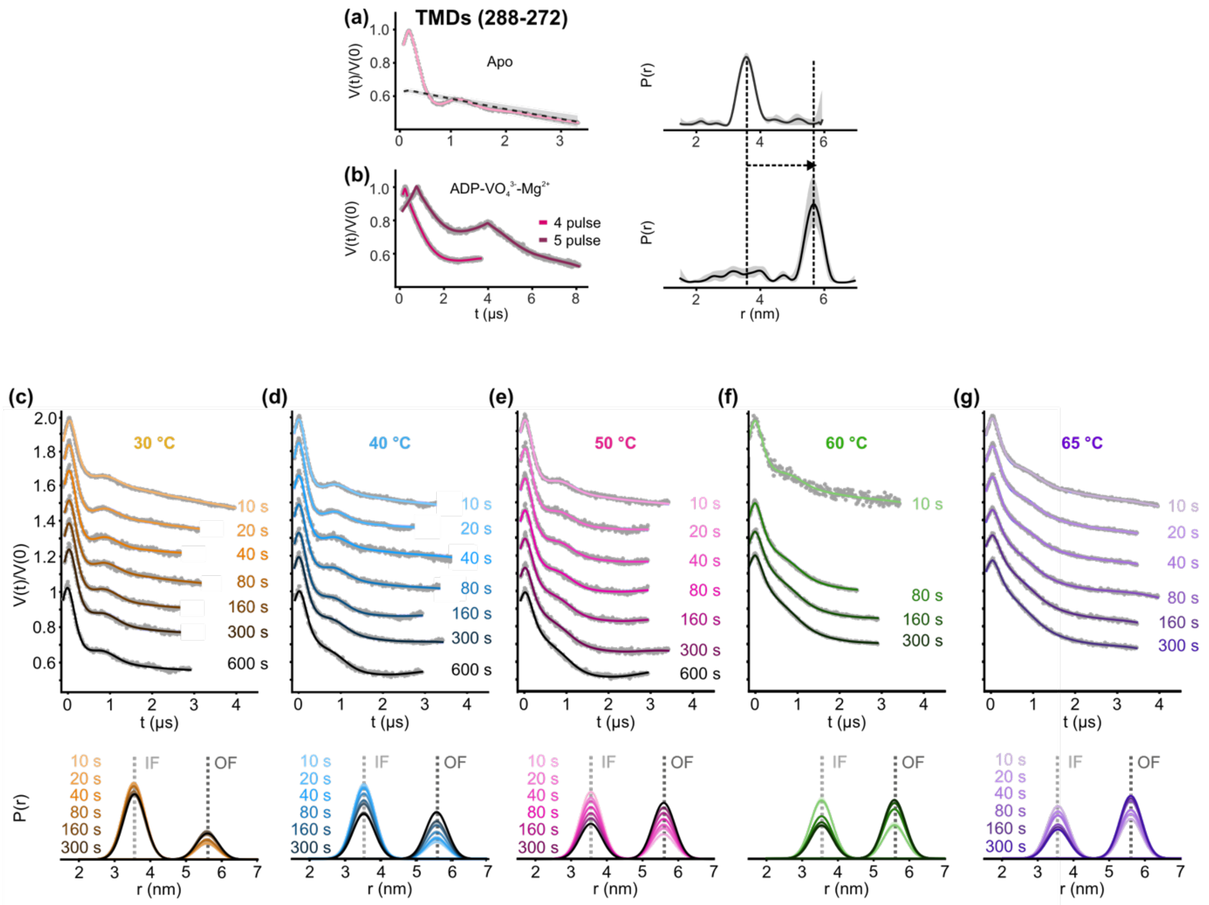


**Supplementary figure 1. Time-resolved PELDOR measurements at the TMDs using the double cysteine variant TmrA 288C – TmrB 272C.** The data corresponds to the results presented in Figure 1e. (**a**) The distances corresponding to the IF (apo) and (**b**) OF (vanadate-trapped, $Mg^{2+}$-ADP-$VO_4^{3-}$) conformations were experimentally determined. For the OF state, both 4pulse- and 5pulse-PELDOR data were measured and globally analyzed. This experimentally determined two-Gaussian model was used to analyze TR-PELDOR data. TmrAB was incubated with ATP-EDTA at (**c**) 30 °C, (**d**) 40 °C, (**e**) 50 °C, (**f**) 60 °C, and (**g**) 65 °C for increasing time intervals as indicated. Primary PELDOR data V(t)/V(0), overlaid with the fit, are shown. Data were globally analyzed using the DeerLab program[5]. The corresponding distances and their probability with a 95% confidence interval (which is invisible if thinner than the line width) are given in the bottom panels.

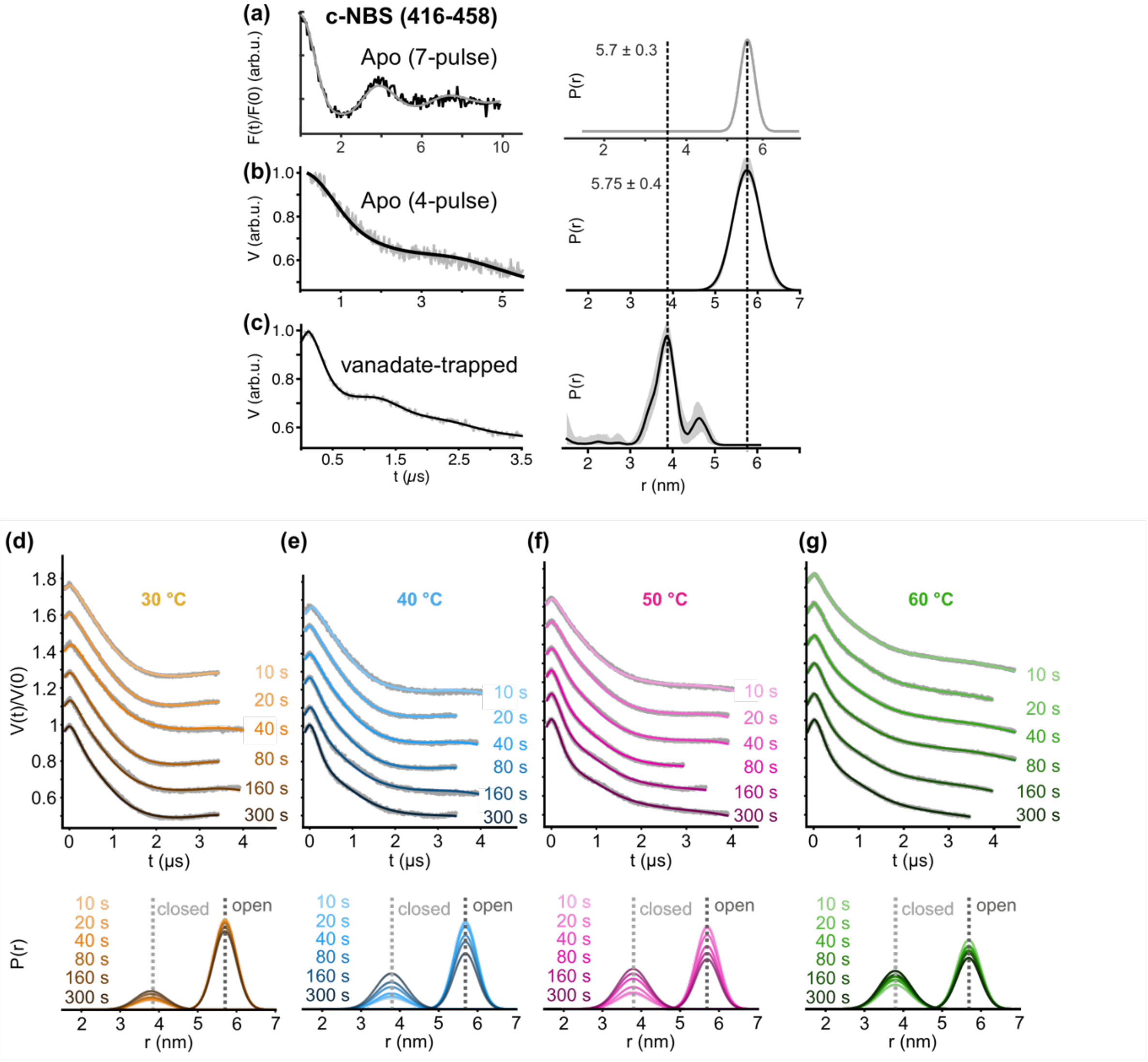


**Supplementary figure 2. TR-PELDOR at the consensus NBS (c-NBS) using the double cysteine variant TmrA 416C – TmrB 458C.** The data corresponds to the results presented in Figure 1f. (**a**) The distances corresponding to the open (apo) and (**b**) closed (vanadate-trapped, $Mg^{2+}$-ADP-$VO_4^{3-}$) conformations were experimentally determined. For the open state, both 4pulse- and 7pulse-PELDOR(2) data were measured, and the longer dipolar evolution for the latter permitted a precise determination of the width of the distance distribution. This experimentally determined two-Gaussian model was used to analyze TR-PELDOR data. TmrAB was incubated with ATP-EDTA at (**c**) 30 °C, (**d**) 40 °C, (**e**) 50 °C, and (**f**) 60 °C for increasing time intervals as indicated. Primary PELDOR data V(t)/V(0), overlaid with the fit, are shown. Data were globally analyzed using the DeerLab program[5]. The corresponding distances and their probability with a 95% confidence interval (which is invisible if thinner than the line width) are given in the bottom panels.

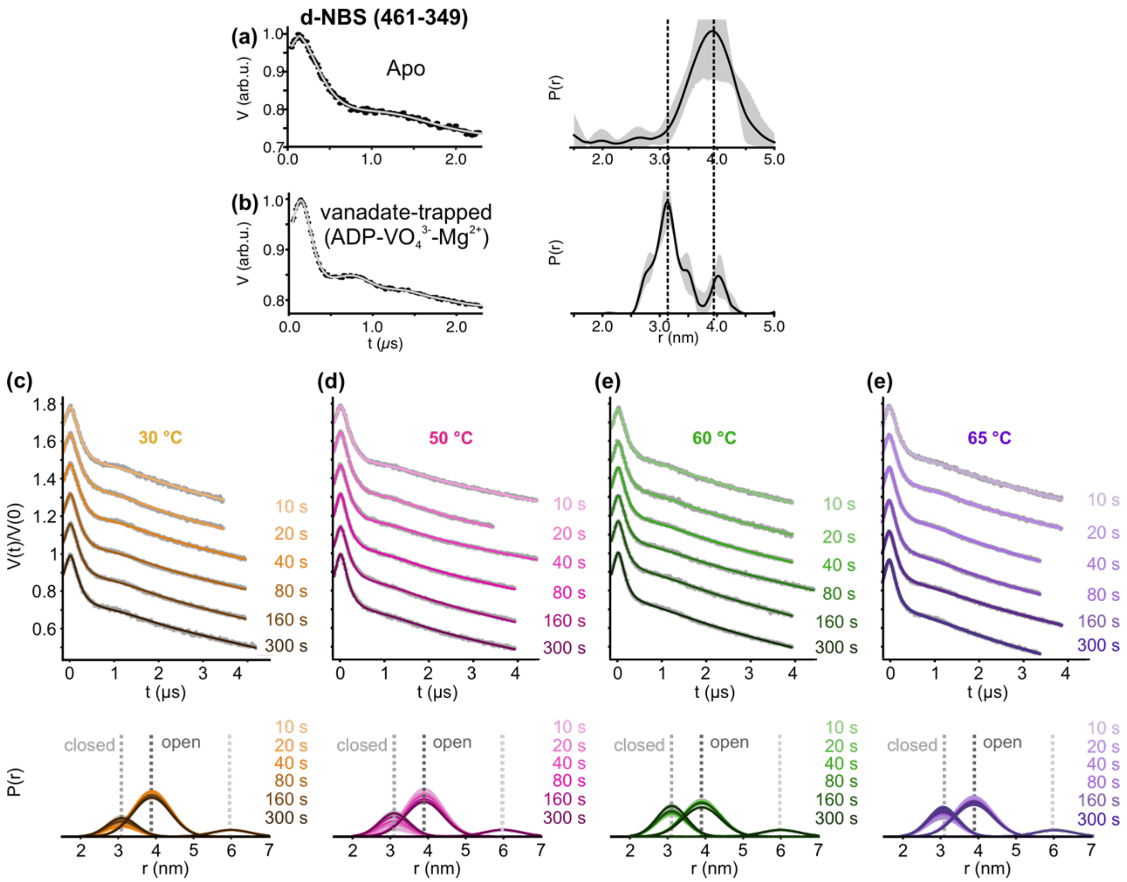


**Supplementary figure 3. TR-PELDOR at the degenerate NBS (d-NBS) using the double cysteine variant TmrA 461C – TmrB 349C.** The data corresponds to the results presented in Figure 1g. (**a**) The distances corresponding to the open (apo) and (**b**) closed (vanadate-trapped, $Mg^{2+}$-ADP-$VO_4^{3-}$) conformations were experimentally determined. This experimentally determined two-Gaussian model was used to analyze TR-PELDOR data. TmrAB was incubated with ATP-EDTA at (**c**) 30 °C, (**d**) 50 °C, (**e**) 60 °C, and (**f**) 65 °C for increasing time intervals as indicated. Primary PELDOR data V(t)/V(0), overlaid with the fit, are shown. Data were globally analyzed using the DeerLab program[5]. The corresponding distances and their probability with a 95% confidence interval (which is invisible if thinner than the line width) are given in the bottom panels. Overall, due to faster kinetics, the equilibrium dynamics cannot be resolved using the experimental setup.

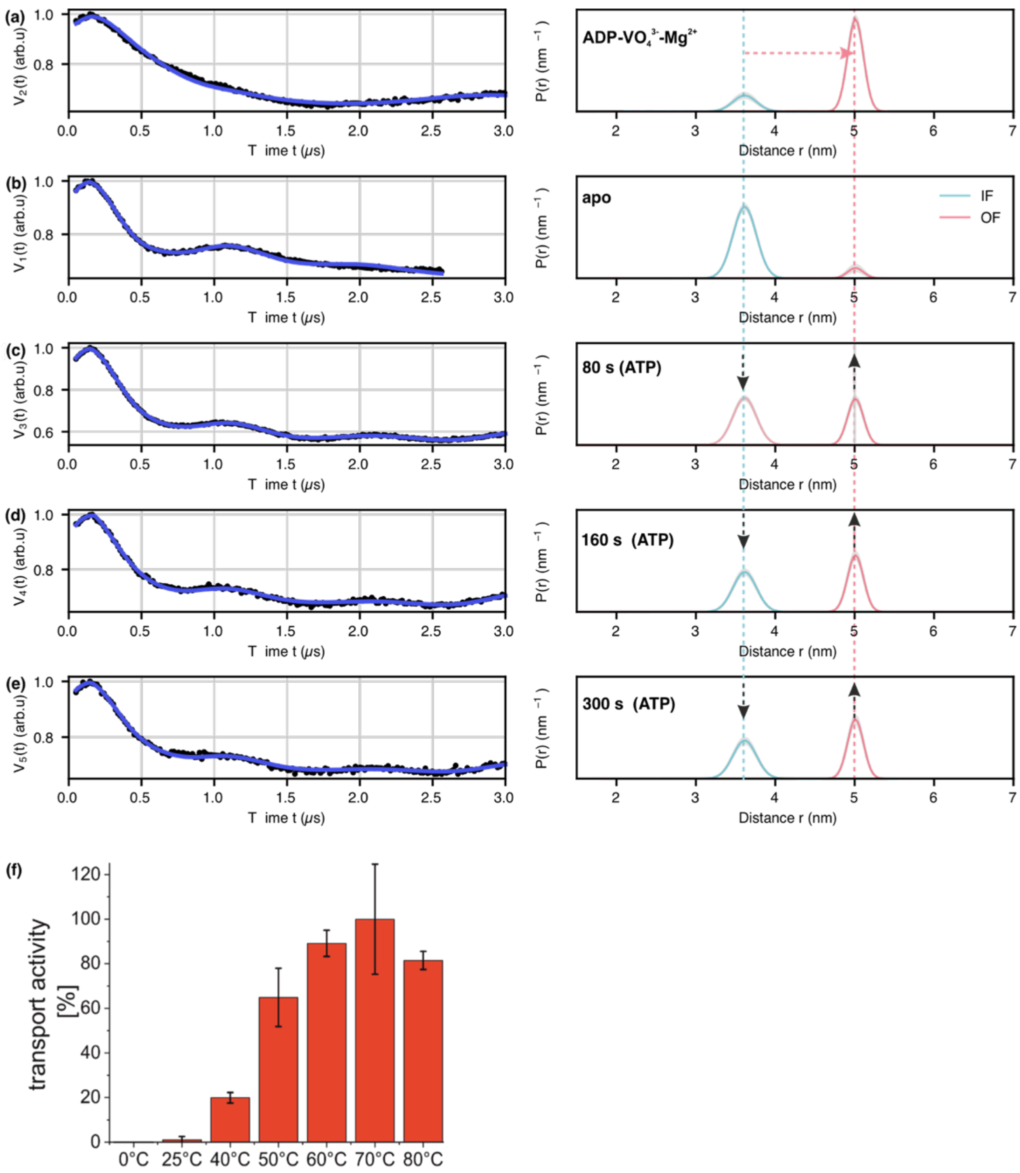


**Supplementary figure 4. Time-resolved PELDOR measurements at the TMDs using the double cysteine variant TmrA 288C – TmrB 272C in nanodiscs.** (**a**) The distances corresponding to the OF (vanadate-trapped, $Mg^{2+}$-ADP-$VO_4^{3-}$) and (**b**) IF (apo) conformations. The data were analyzed according to the experimentally determined two-Gaussian model as shown in Supplementary figure 1. (**c-e**) TmrAB was incubated with ATP-EDTA at 40 °C, and samples were collected at (**c**) 80 s, (**d**) 160 s, and (**e**) 300 s. Primary PELDOR data V(t)/V(0), overlaid with the fit, are shown. Data were globally analyzed using the two-Gaussian model employing the DeerLab program[5]. The corresponding distances and their probability with a 95% confidence interval (which is invisible if thinner than the line width) are given in the right panels. The data show that the transition reached the equilibrium already at the first sampling point (80 s) and minimally changed afterwards, revealing a faster kinetics in comparison to the corresponding micellar sample (Supplementary figure 1d). (**f**) Temperature-dependent transport of a peptide substrate by liposome-reconstituted wild-type TmrAB. Error bars indicate standard deviation (*n* = 3).

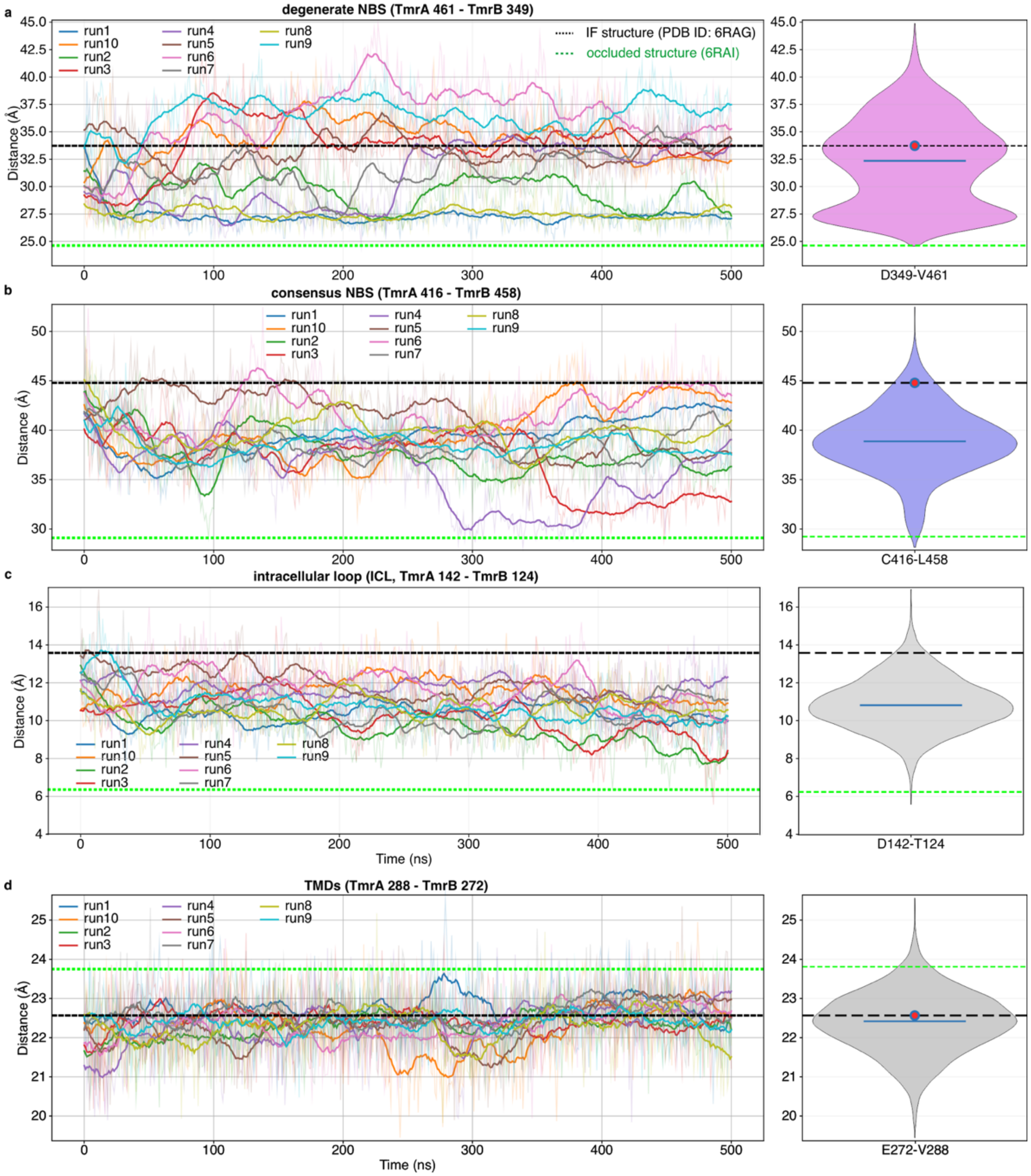


**Supplementary figure 5-I. Time evolution and pooled distance distributions in the inward-facing TmrAB** (E → Q variant) **with bound ATP–$Mg^{2+}$ during IF to OCC transition.** The data corresponds to the analysis presented in Figure 2a (IF to OCC part of the unbiased MD simulations). The distances monitored are (**a**) V461-D349 (d-NBS), (**b**) C416–L458 (c-NBS), and (**d**) V288–E272 (TMDs), as established by PELDOR experiments (Supplementary figures 1-3), and (**c**) D142–T124 (the intracellular loop, ICL, highlighted as spheres in Supplementary figure 6f). During the inward-facing (IF) to the occluded (OCC) transition, the NBSs and the ICL close, and the TMDs slightly open (see Figure 1a). In the left panels, each colored trace represents one independent trajectory, with faint lines showing the raw data and bold lines showing the smoothed trajectories. In the right panels, violin plots show the pooled distribution **(**from three minimum heavy-atom distances**)** of distances across all trajectories (10 x 500 ns simulations); the blue horizontal line marks the median, and the red circle marks the mean value at the starting frame. For structural comparison, the distance corresponding to the inward-facing reference structure is indicated by a black dashed line and the occluded-state reference structure by a green dotted line in both the time-series and violin panels. Together, these comparisons show how the simulated inward-facing ensemble samples conformations relative to the inward-facing and occluded structural states. An independent PCA analysis of the data is presented in Supplementary figure 5-IIIa.

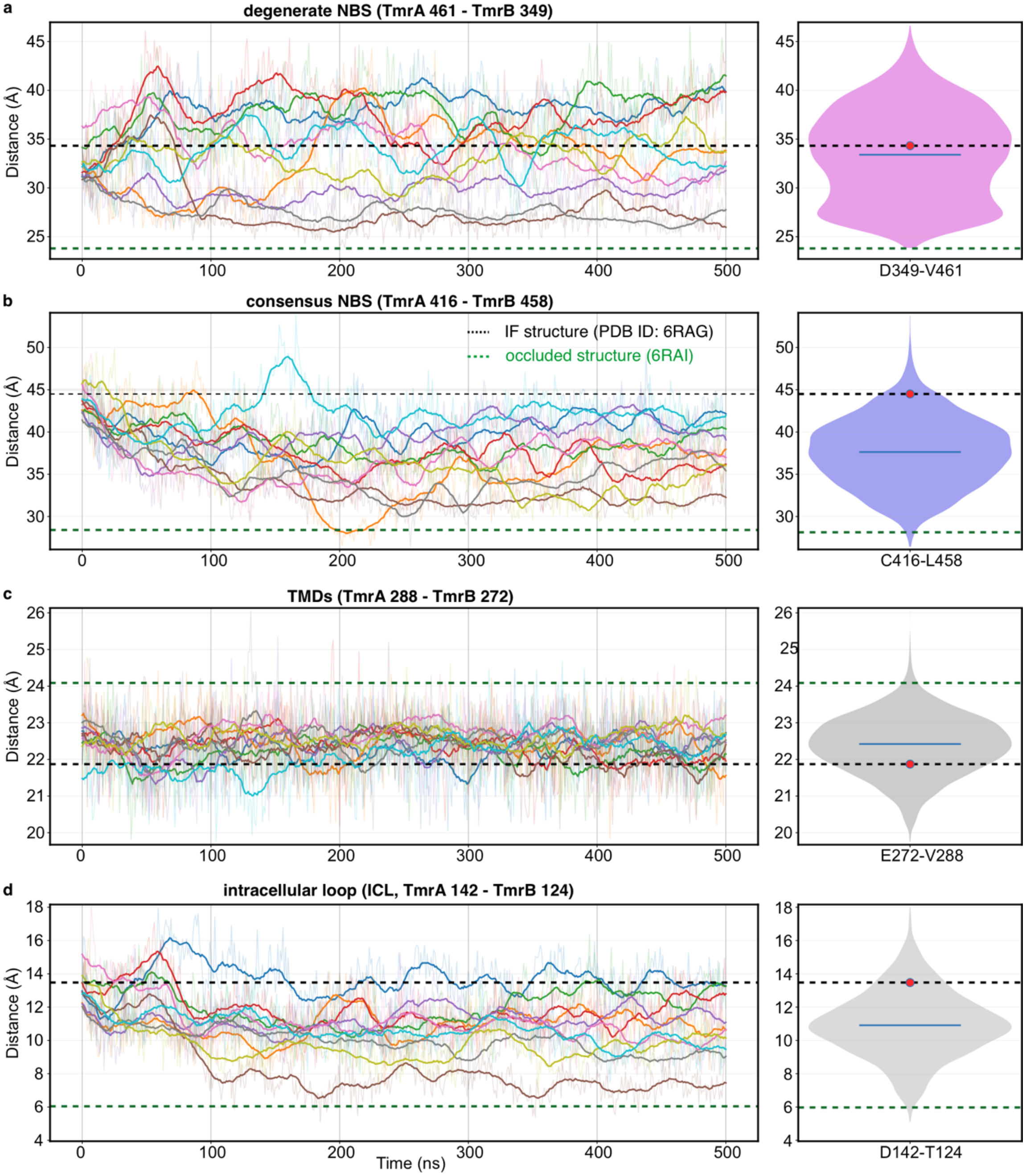


**Supplementary figure 5-II. Time evolution and pooled distributions in the inward-facing TmrAB (WT) with bound ATP–Mg²⁺ during IF to OCC transition.** The distances monitored are (**a**) V461-D349 (d-NBS), (**b**) C416–L458 (c-NBS), and (**d**) V288–E272 (TMDs), as established by PELDOR experiments (Supplementary figures 1-3), and (**c**) D142–T124 (the intracellular loop, ICL, highlighted as spheres in Supplementary figure 6f). In the right panels, violin plots show the pooled distribution (from three minimum heavy-atom distances) of distances across all trajectories (10 x 500 ns simulations). Panel descriptions are identical to those given for the E→Q variant in Supplementary figure 5-I. The corresponding PCA analysis is presented in Supplementary figure 5-III.

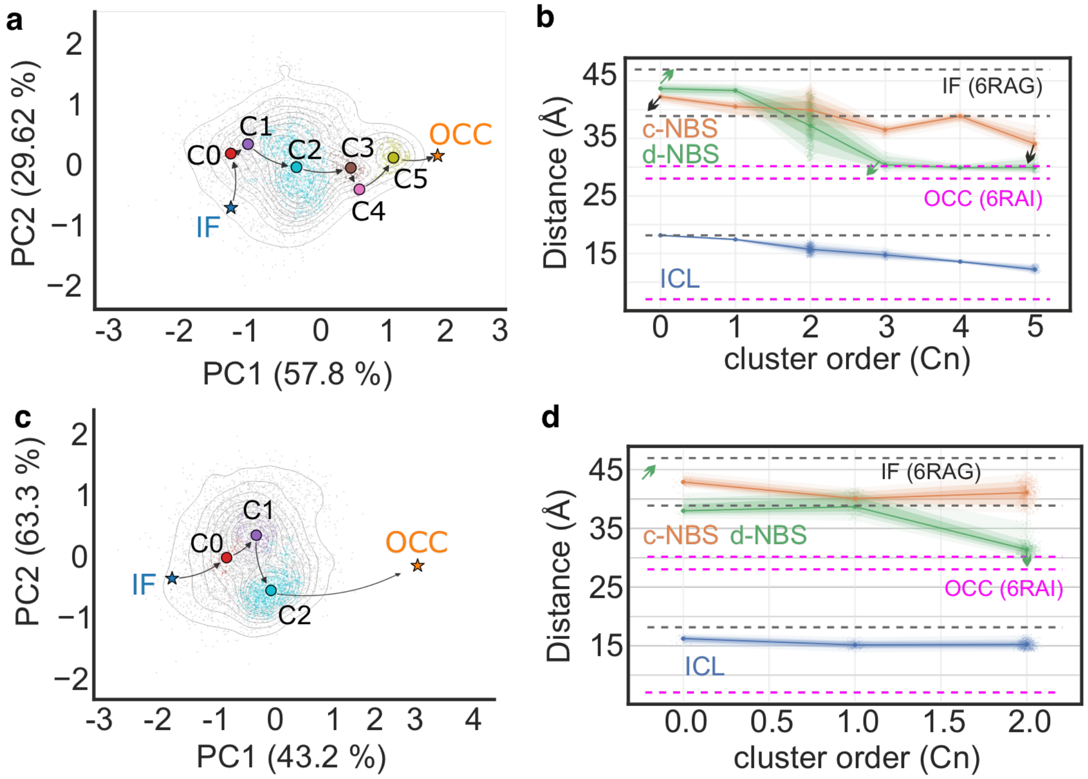


**Supplementary figure 5-III. Conformational sampling from unbiased MD simulations for the IF→OCC transition in TmrAB (WT and E→Q variant). (a-b)** The images correspond to the IF to OCC transition part of the PCA analysis, as presented in Figure 2a for the E→Q variant (corresponding trajectories are presented in Supplementary figure 5-I) and (**c-d**) the data for the WT variant (the corresponding trajectories in Supplementary figure 5-II). Three $C_\alpha - C_\alpha$ distances (D142–T124 (ICL), C416–L458 (c-NBS), and D349–V461 (d-NBS)) as established by PELDOR experiments (Supplementary figures 1-3) were z-scored and equally weighted (⅓ each) as the features for the principal component analysis (10 x 500 ns simulations). Points are colored by HDBSCAN cluster; gray dots are noise. Centroids $C_n$ are ordered along the IF→OCC axis in the full feature space; the arrow chain links IF (blue star) through centroids to OCC structure (orange star), and the corresponding distances are shown in the right panels. The reference distances corresponding to the IF (6RAG) and OCC (6RAI) structures are indicated with grey and magenta dotted lines in panels b and d, respectively.

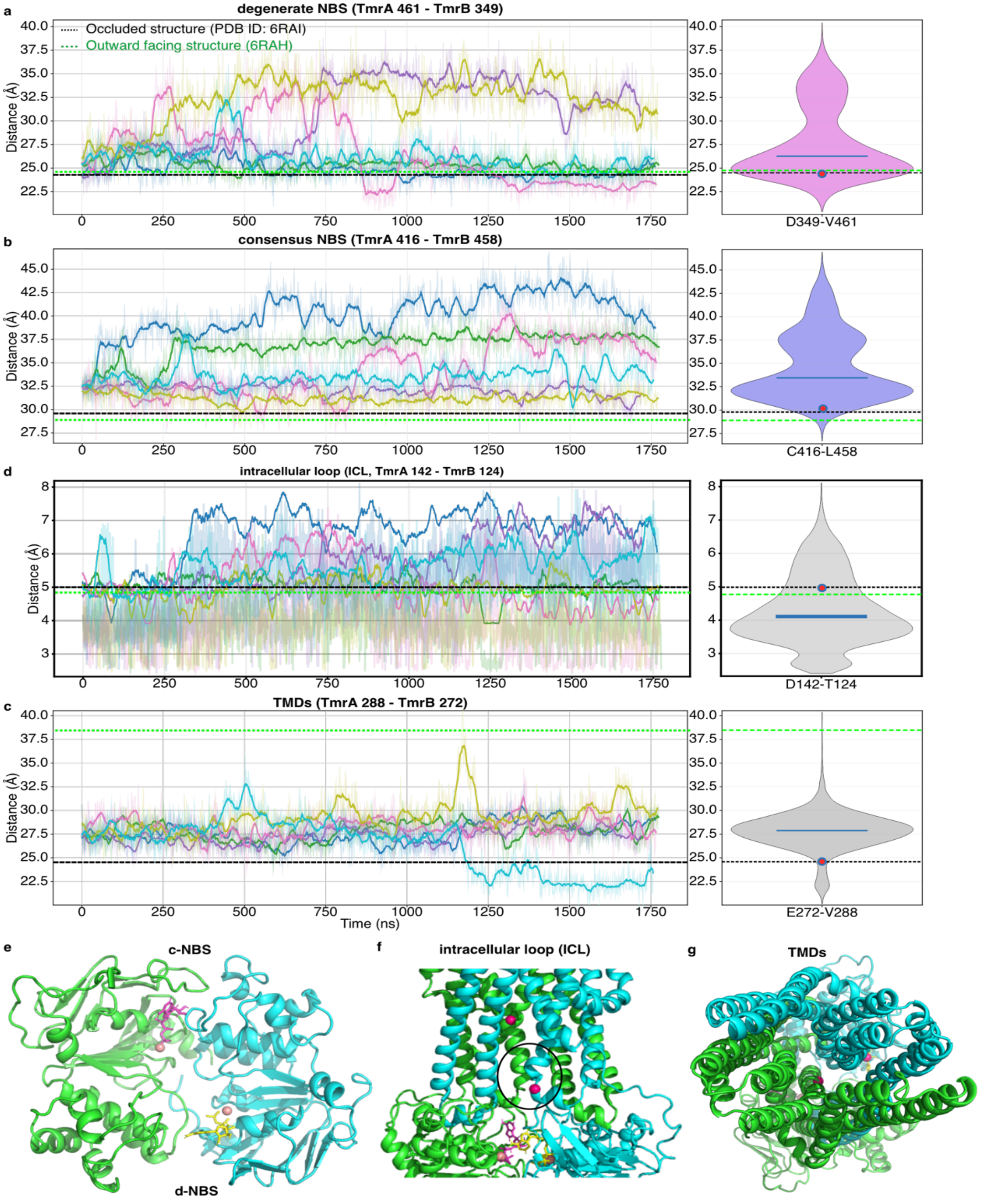


**Supplementary figure 6. Time evolution and pooled distributions in TmrAB (E→Q variant) with bound ATP–$Mg^{2+}$ during OCC→OF transition**. The trajectories correspond to the OCC→OF part of the PCA analysis as presented in Figure 2a. The distances monitored are (**a**) V461-D349 (d-NBS), (**b**) C416–L458 (c-NBS), and (**d**) V288–E272 (TMDs), as established by PELDOR experiments (Supplementary figures 1-3), and (**c**) D142–T124 (the intracellular loop, ICL, f). During the occluded (OCC) to outward-facing (OF) transition observed here, the two NBSs and the ICL remain closed, and the TMDs open (see Figure 1a). In the left panels, each colored trace represents one independent trajectory (6 x 1.7 μs simulations), with faint lines showing the raw data and bold lines showing the smoothed trajectories. In the right panels, violin plots show the pooled distribution of (three minimum heavy-atom) distances across all trajectories; the blue horizontal line marks the median, and the red circle marks the mean value at the starting frame. For structural comparison, the occluded-state reference structure is indicated by a black dashed line and the outward-facing reference structure by a green dotted line in both the time-series and violin panels. In agreement with a lower barrier, the d-NBS exhibited independent flexibility, also revealing an open conformation (e) when the c-NBS (e) and ICL (f) are closed (highlighted inside the circle), and the TMDs are open and outward-facing (g). In panel f, positions D142 (at top) and T124 (at bottom), used to monitor the ICL, are highlighted as magenta spheres. Together, these comparisons show how the simulated occluded-state ensemble samples conformations relative to the occluded and outward-facing reference states.

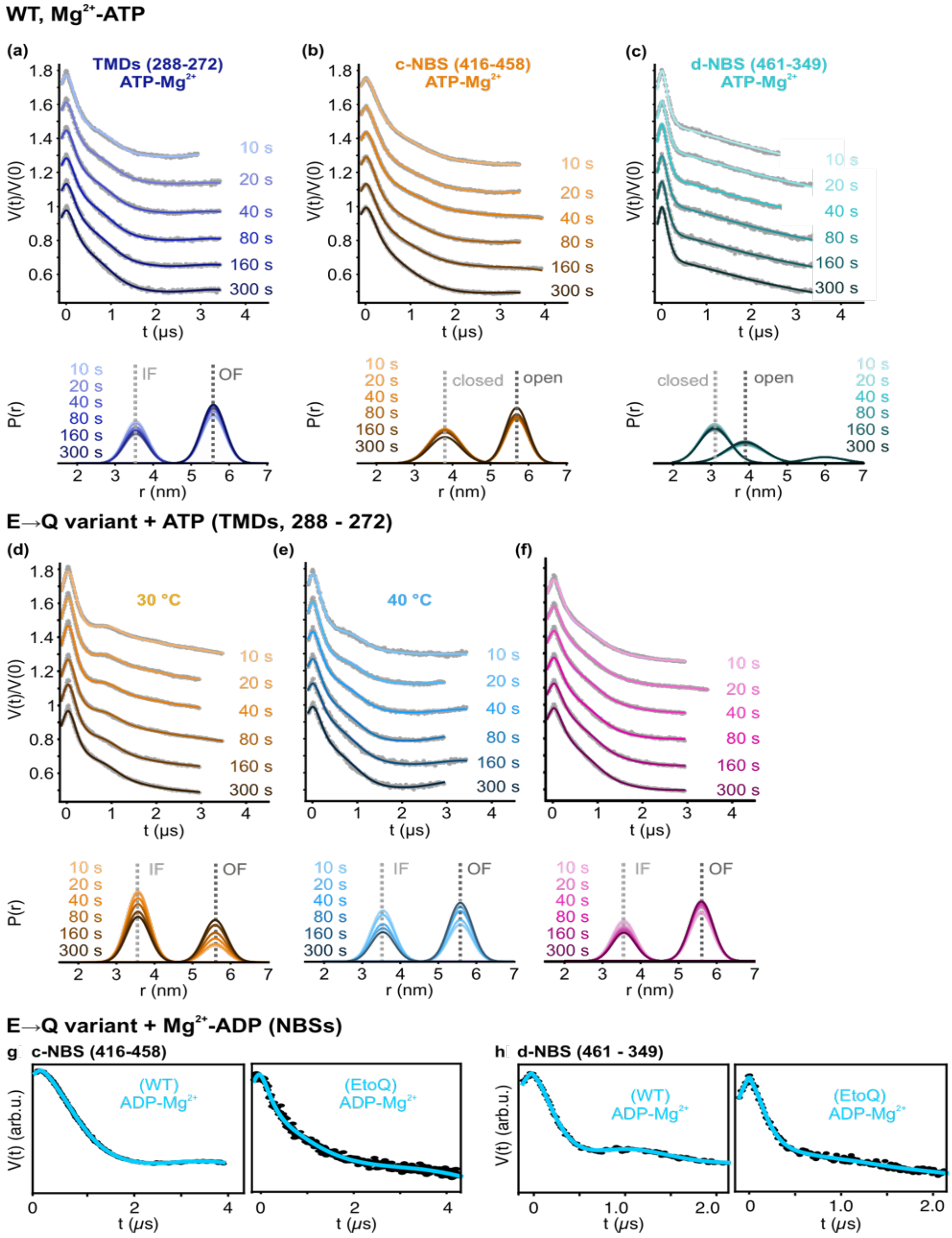


**Supplementary figure 7. PELDOR experiments at the TMDs and the NBSs in the WT ($Mg^{2+}$-ATP) or the E→Q variant (ATP or $Mg^{2+}$-ADP).** TmrAB was incubated with $Mg^{2+}$-ATP at 40 °C, and TR PELDOR experiments were performed as indicated for (**a**) TMDs, (**b**) c-NBS, and (**c**) the d-NBS (corresponding to Figure 2b-c). (**d-f**) For the E→Q variant, the IF↔OF transition was observed (40 °C, ATP alone) at the TMDs in a time-resolved manner at different temperatures, as indicated (kinetics presented in Figure 2d). The data sets were globally analyzed using an experimentally determined two-Gaussian model (Supplementary figures 1-3). Primary PELDOR data V(t)/V(0), overlaid with the fit, are shown and analyzed using the DeerLab program[5]. The corresponding distances and their probability with a 95% confidence interval (which is invisible if thinner than the line width) are given below. Overall, due to faster kinetics in the presence of $Mg^{2+}$- or with the E→Q substitution, the equilibrium dynamics cannot be time-resolved. (**g-h**) Primary PELDOR data corresponding to the WT or the E→Q variant at the c-NBS and the d-NBS as indicated in the presence of $Mg^{2+}$-ADP. The corresponding distance distributions (using a two-Gaussian fit, Supplementary figure 2-3) are presented in Figure 3d-e.

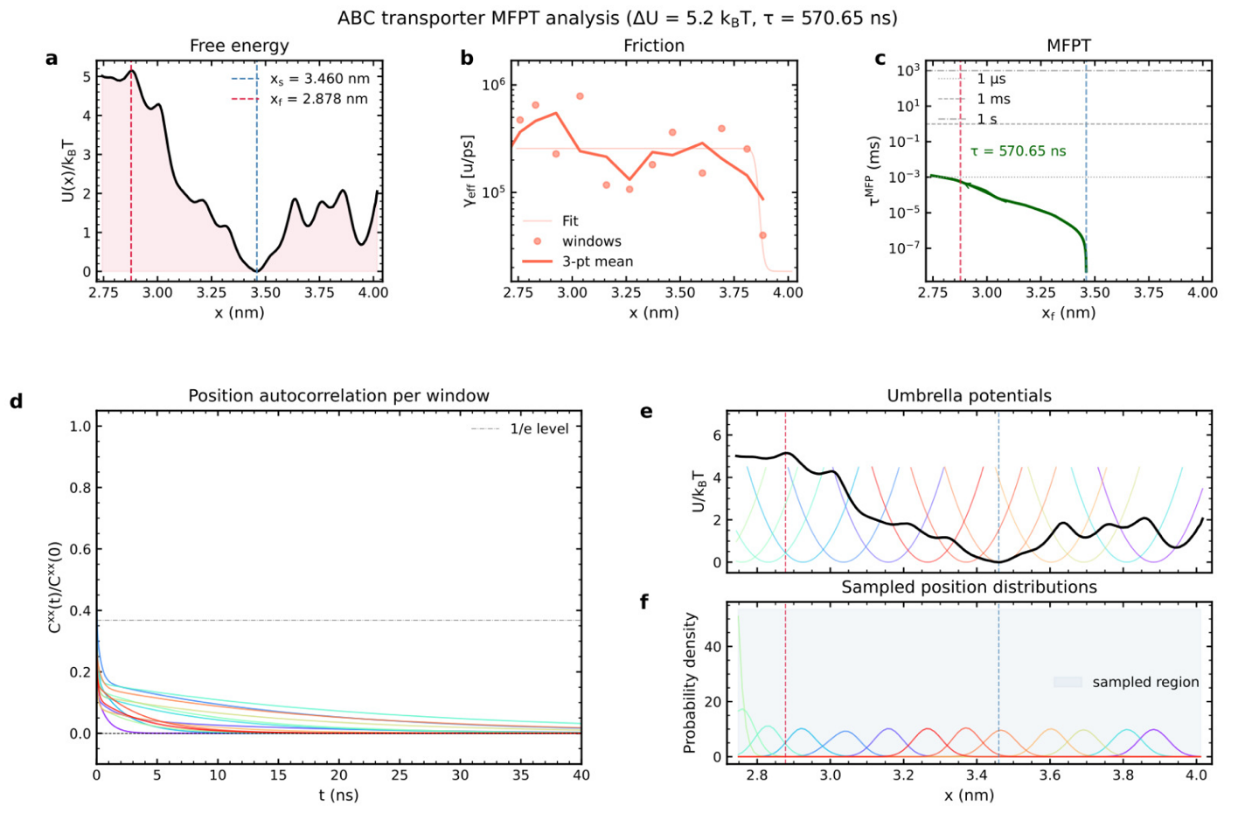


**Supplementary figure 8-I. MFPT analysis of the degenerate nucleotide-binding site** (d-NBS, TmrA V461 – TmrB D349)**.** This reaction coordinate was established from PELDOR spectroscopy (Supplementary figure 3). The d-NBS site is described by an independent distance coordinate, which is kinetically uncoupled from c-NBS (as observed from PELDOR data, Figure 1 f–g). (**a**) Free-energy profile U(x)/$k_B$T obtained from umbrella sampling and weighted histogram analysis (WHAM), with the basin starting point $x_s$, (blue dashed line) and barrier $x_f$ (red dashed line) marked, giving $\Delta U$ = 4.1 $k_B$T for this site. The reaction coordinate is defined as the distance between the centers of mass of residue groups 461 and 349. (**b**) Position-dependent effective friction $\gamma_{eff}(x)$ computed from the position autocorrelation function (**e**), individual umbrella windows (red circles), 3-point geometric-mean trend (bold red line), and sigmoid fit (faint red line) used in the Fokker–Planck integration are shown. (**c**) Position-dependent diffusion coefficient $D_{eff}(x) = k_BT / \gamma_{eff}(x)$, shown in the same style as panel B. (**d**) Mean first-passage time (MFPT, denoted as $\tau^{MFP}$) as a function of the final position $x_f$, computed by numerical integration of the one-dimensional Fokker–Planck equation. The MFPT for the full barrier crossing of the d-NBS is 285.32 ns. (**e**) Normalized position autocorrelation function $C^{xx}(t)/C^{xx}(0)$ for each umbrella window (colored by window position), and the dashed line marks the 1/*e* level. (**f**) PMF (black) overlaid with the harmonic umbrella biasing potentials of all 13 windows (colored), illustrating the umbrella coverage along the reaction coordinate. (**g**) Probability density of the sampled positions in each umbrella window; the shaded region marks the full sampled range. Simulation details: 13 umbrella windows along the reaction coordinate, harmonic spring constant $k$ = 1000 kJ mol$^{-1}$ nm$^{-2}$. Each window was simulated for 500 ns.

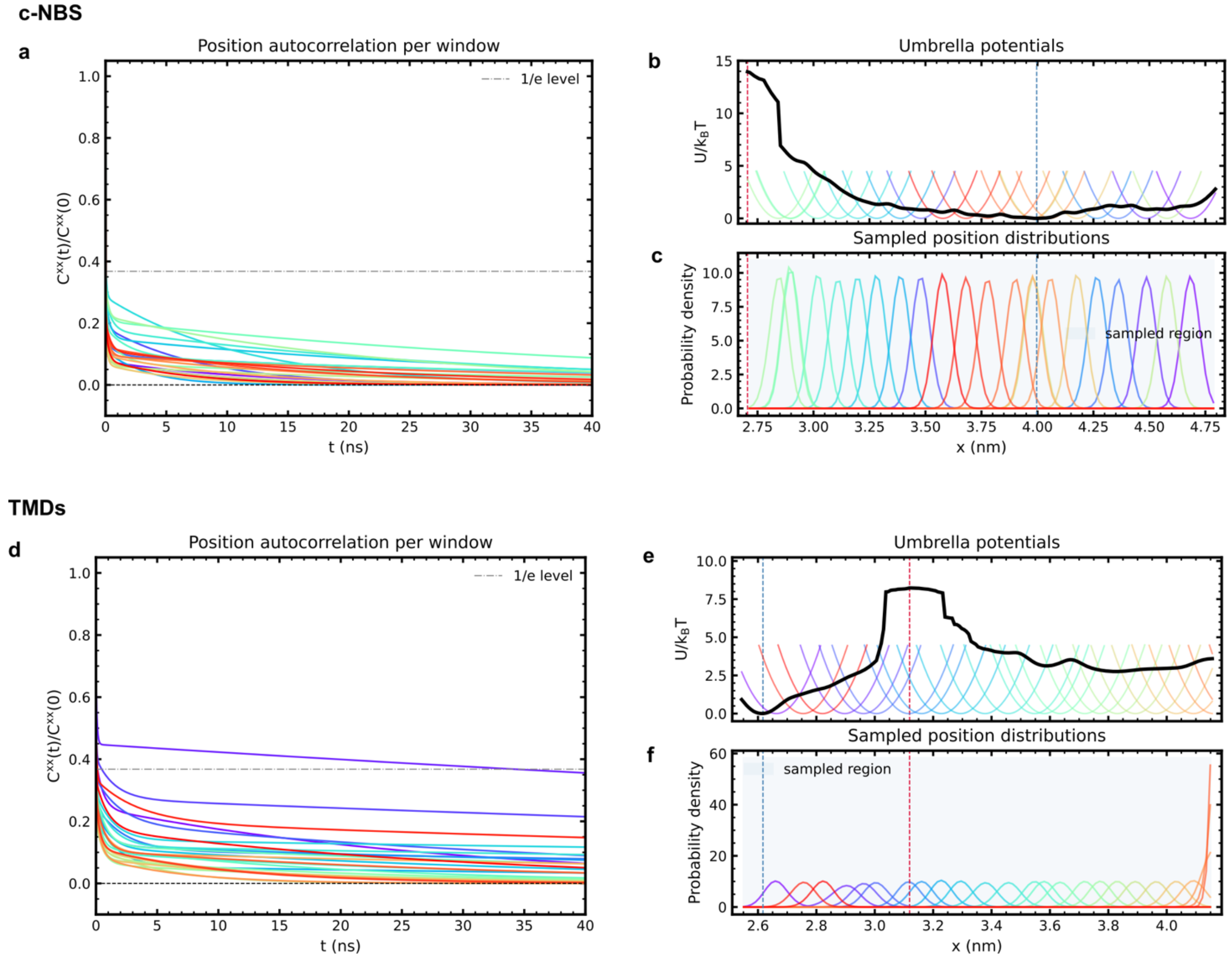


**Supplementary figure 8-II. Probability density distributions of the sampled positions in each umbrella-sampling window**. The collective variables, consensus nucleotide-binding site (c-NBS; C416–L458) and the transmembrane domains (TMDs; V288–E272) in TmrAB are established from PELDOR spectroscopy (Supplementary figure 1-2). (**a,d**) Normalized position autocorrelation function $C^{xx}(t)/C^{xx}(0)$ for each umbrella window (colored by window position), and the dashed line marks the 1/*e* level. (**b,e**) PMF (black) overlaid with the harmonic umbrella biasing potentials of all 13 windows (colored), illustrating the umbrella coverage along the reaction coordinate. (**c,f**) Probability density of the sampled positions in each umbrella window; the shaded region marks the full sampled range. The reaction coordinate was defined as the distance between the centers of mass of the selected residue pairs. Simulation details: 13 umbrella windows along the reaction coordinate, harmonic spring constant $k$ = 1000 kJ mol$^{-1}$ nm$^{-2}$. Each window was simulated for 500 ns. The blue dashed line indicates the position of the minimum-energy state, whereas the red dashed line marks the location of the energy barrier.

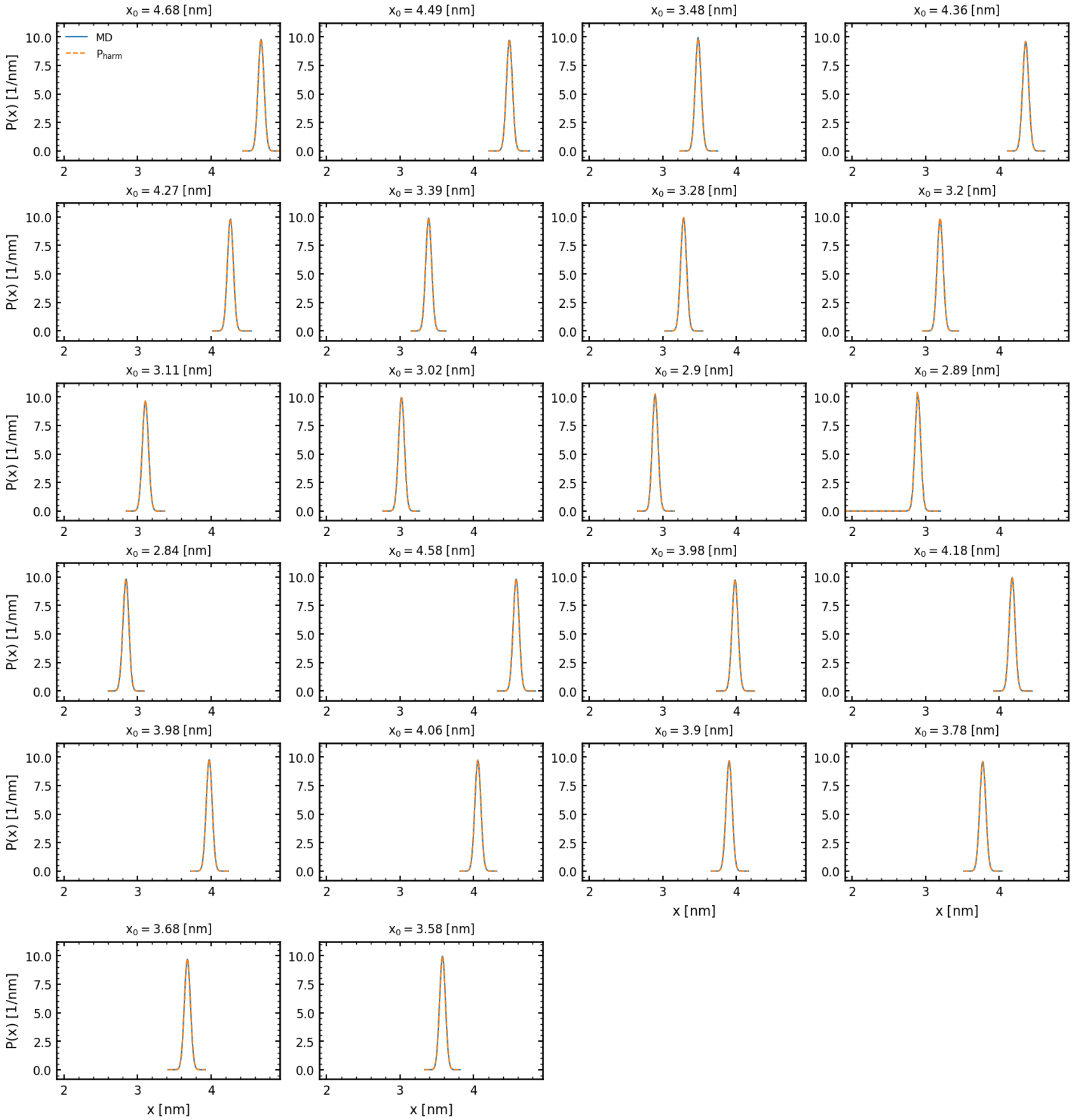


**Supplementary figure 9. Harmonic approximation of c–NBS umbrella sampling distributions.** Probability distributions P(x) of the c–NBS (C416 – L458) collective variable x obtained from individual umbrella sampling simulations. Blue curves show histograms from the molecular dynamics trajectories, while dashed orange curves show the corresponding harmonic reference distributions, $P_{harm}(x)$, expected from the applied umbrella potential. The close agreement between P(x) and $P_{harm}(x)$ across all umbrella windows supports the local harmonic approximation used for the friction analysis. The titles denote $x_0$, the center of the restraining potentials along the c–NBS collective variable.

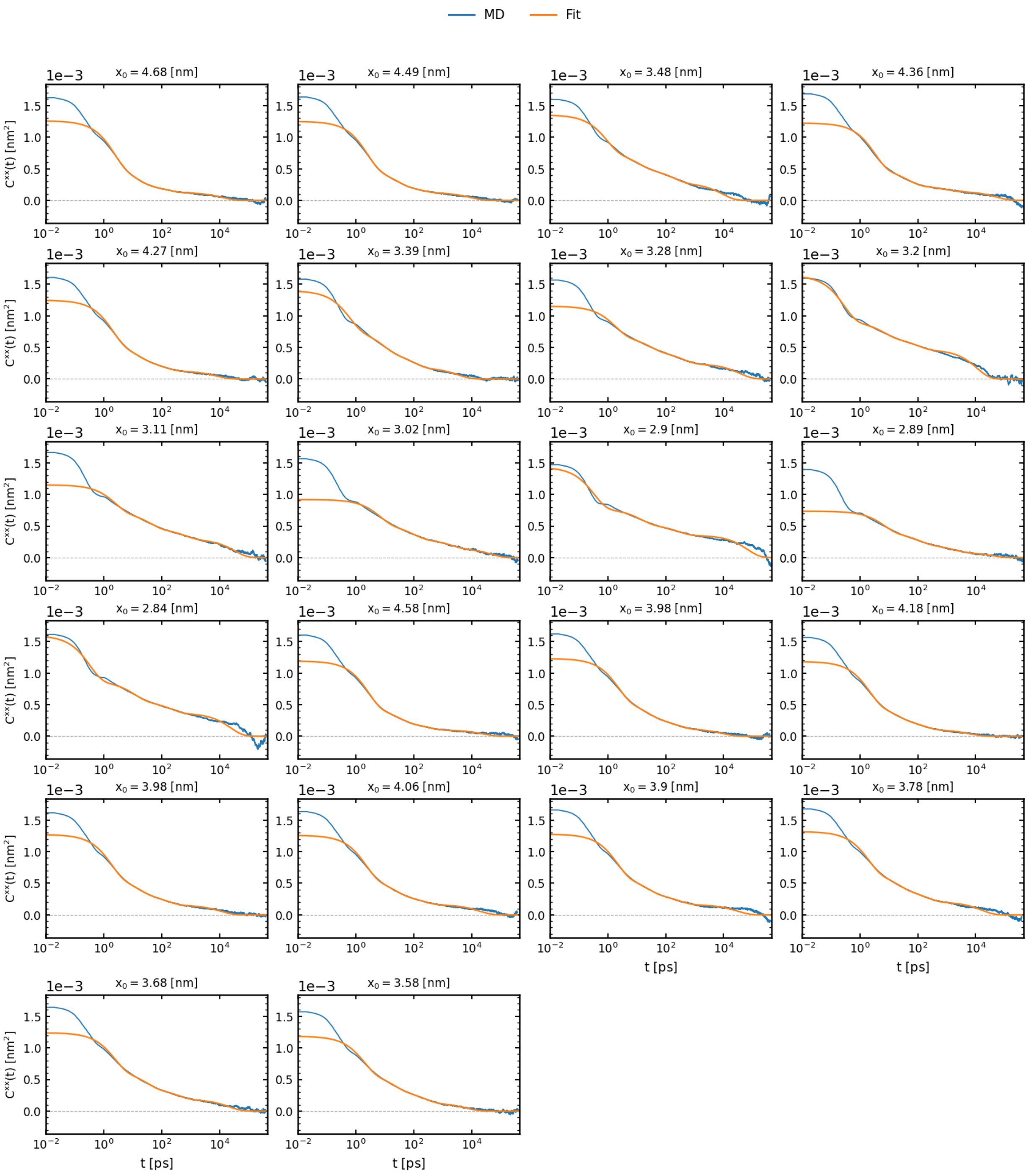


**Supplementary figure 10. Autocorrelation functions of the c–NBS collective variable.** Position autocorrelation functions of the c–NBS (C416 – L458) collective variable,

$$C_{xx}(t) = \langle (x(0) - \langle x \rangle)(x(t) - \langle x \rangle) \rangle,$$

computed from the umbrella sampling trajectories. Blue curves show the simulation data, and orange curves show multi-exponential fits used to evaluate the correlation-time integrals. The fits capture the overall relaxation behavior of the C–NBS coordinate in each umbrella window, while deviations at short times and noisy long-time tails reflect inertial effects and finite sampling. The titles denote $x_0$, the centers of the restraining potentials along the C–NBS collective variable.

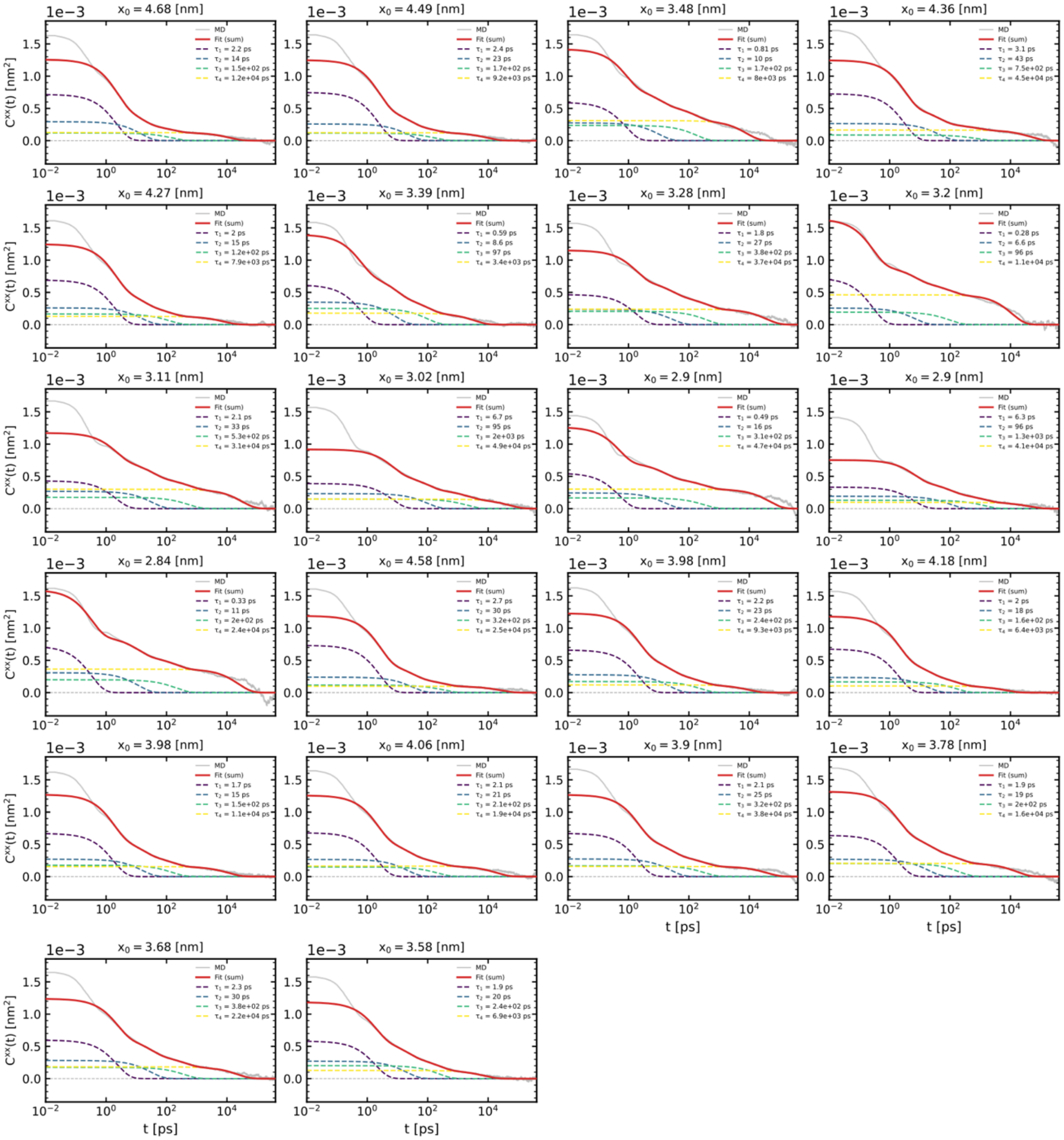


**Supplementary figure 11. Multi-exponential decomposition of the c–NBS position autocorrelation function.** Position autocorrelation functions $C^{xx}(t) = \langle (x(0) - \langle x \rangle)(x(t) - \langle x \rangle) \rangle$ of the c–NBS (C416–L458) collective variable for each umbrella-sampling window (grey, MD), overlaid with the multi-exponential fit (red, sum) and its individual exponential components (dashed), $(g_i/\tau_i)\cdot\exp(-t/\tau_i)$. The component decay times $\tau_i$ are listed in each legend and span from a few ps to ~$10^4$ ps, showing that $C^{xx}(t)$ is governed by several well-separated relaxation timescales. This multi-exponential (memory) structure distinguishes the memory-corrected effective friction $\gamma_{eff}$ (Eq. 4, weighting the squared autocorrelation) from the single-relaxation estimate $\gamma_{pos}$ (Eq. 8, weighting the linear autocorrelation); the two would coincide for a single-exponential $C^{xx}(t)$. Panel titles denote $x_0$, the centre of the umbrella restraining potential along the c–NBS coordinate. Fits use $n_{exp} = 4$.

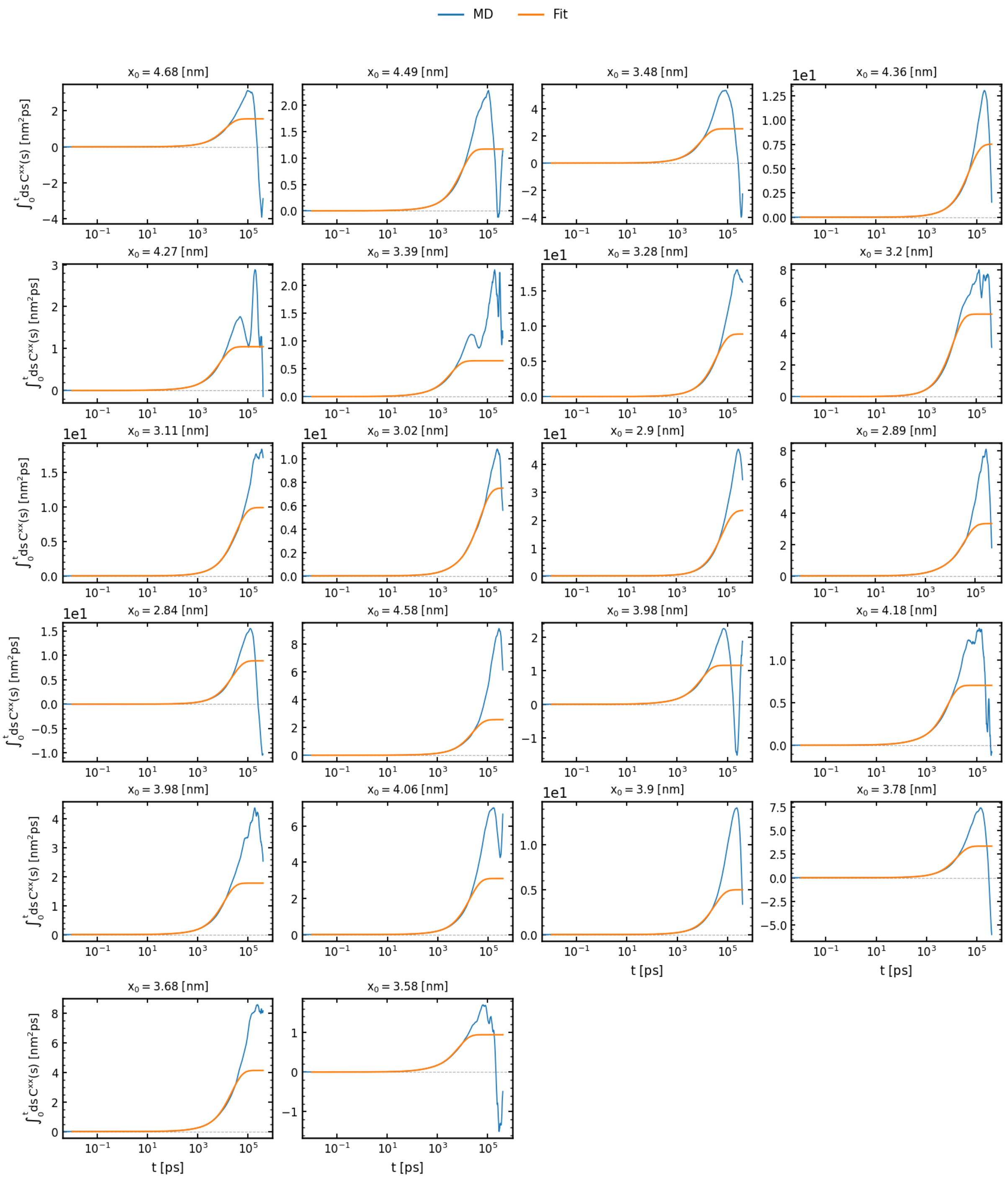


**Supplementary figure 12. Running integrals of the C–NBS autocorrelation functions**. Running integrals $\int_0^t ds\, C_{xx}(s)$ of the autocorrelation functions of the c–NBS (C416 – L458) collective variable. Blue curves show numerical integrals from the simulation data, and orange curves show the corresponding integrals obtained from the fitted autocorrelation functions. The fitted integrals provide smoothed estimates of the long-time correlation contribution used to compute the position-based friction coefficient for motion along the c–NBS collective variable. The titles denote $x_0$, the centers of the restraining potentials.

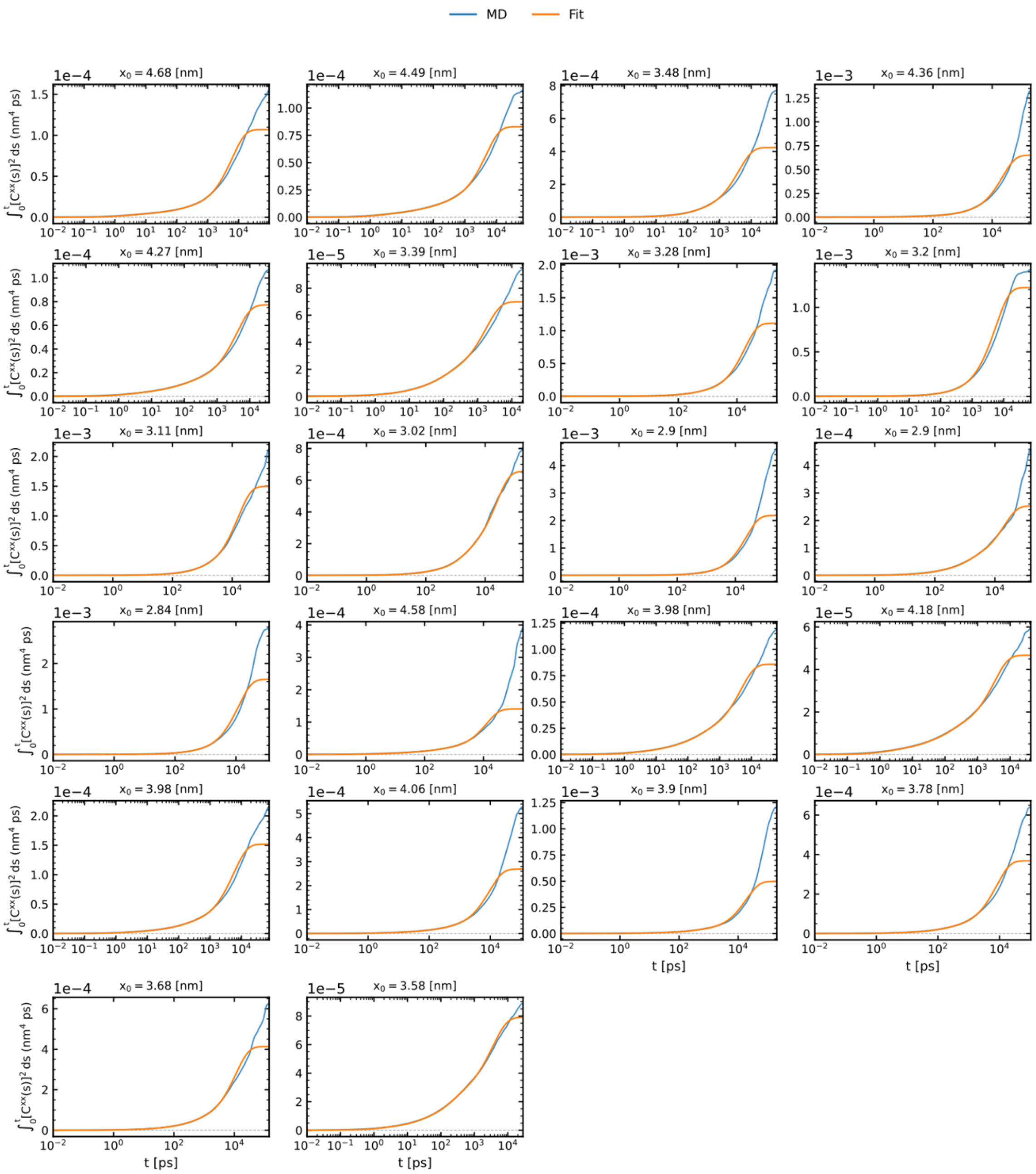


**Supplementary figure 13. Convergence of the squared-autocorrelation integral that determines $\gamma_{eff}$.** Running integrals $\int_0^t [C^{xx}(s)]^2\,ds$ of the c–NBS (C416–L458) collective variable — the quantity entering the effective-friction estimate $\gamma_{eff}$ (Eq. 4) — evaluated directly from the MD autocorrelation (blue) and from its multi-exponential fit (orange). Both reach a common plateau, confirming convergence of the integral that determines $\gamma_{eff}$. Each curve is shown up to the first zero crossing of $C^{xx}(t)$, the lag beyond which the autocorrelation has decayed into the statistical noise floor; this is the same range over which the integral coverage of 0.94–1.00 is reported. Panel titles denote $x_0$, the centre of the umbrella restraining potential along the c–NBS coordinate.

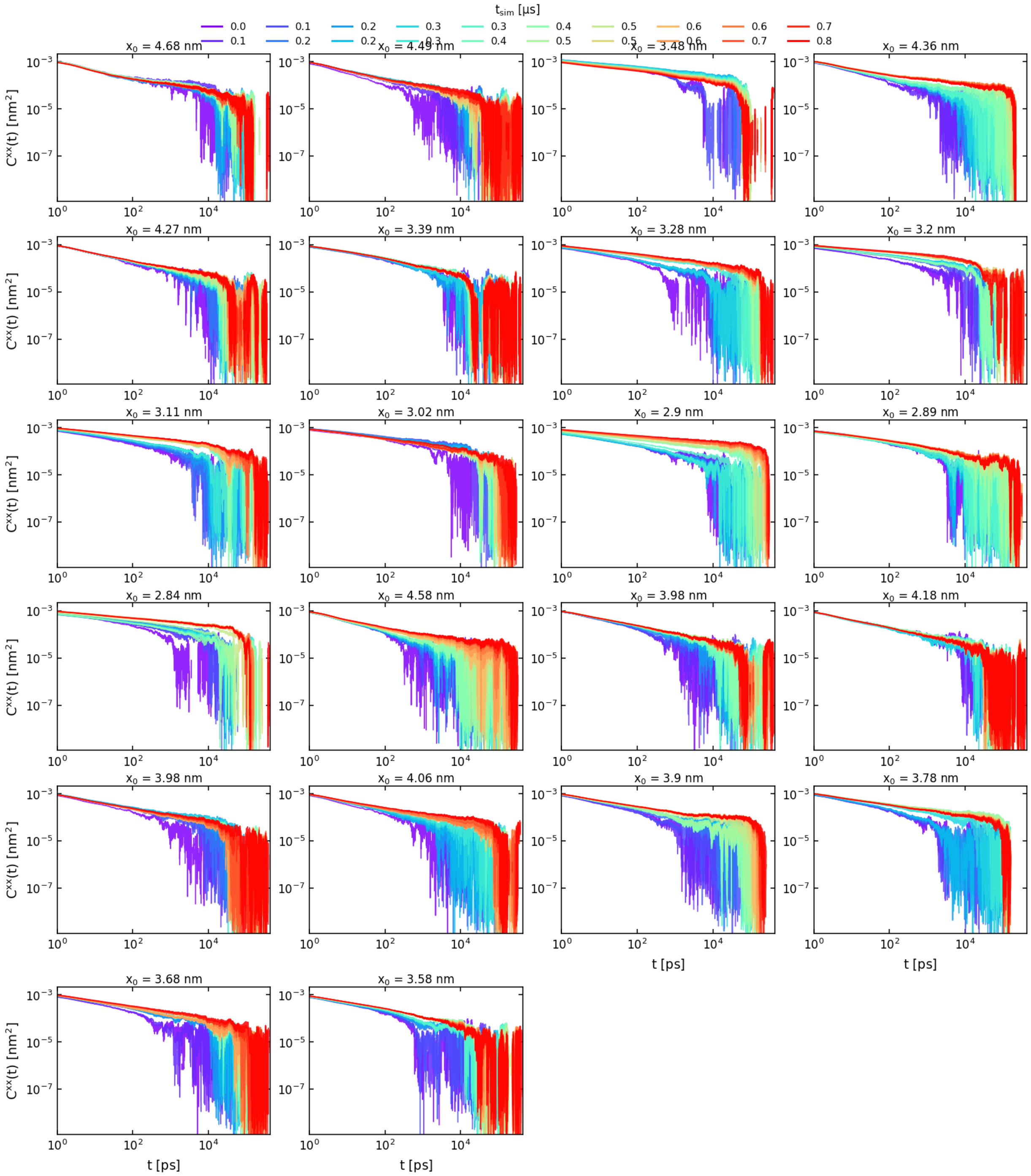


**Supplementary figure 14. Convergence of c–NBS autocorrelation functions.** Autocorrelation functions $C_{xx}(t)$ of the c–NBS (C416 – L458) collective variable computed using increasing amounts of simulation data from the umbrella sampling trajectories. Colors indicate the cumulative simulation time included in the analysis. The overlap of curves at short and intermediate lag times indicates that the dominant relaxation behavior along the c–NBS coordinate is reasonably stable with simulation length, whereas deviations at long lag times reflect reduced statistical sampling. The titles denote $x_0$, the centers of the restraining potentials along the c–NBS collective variable.

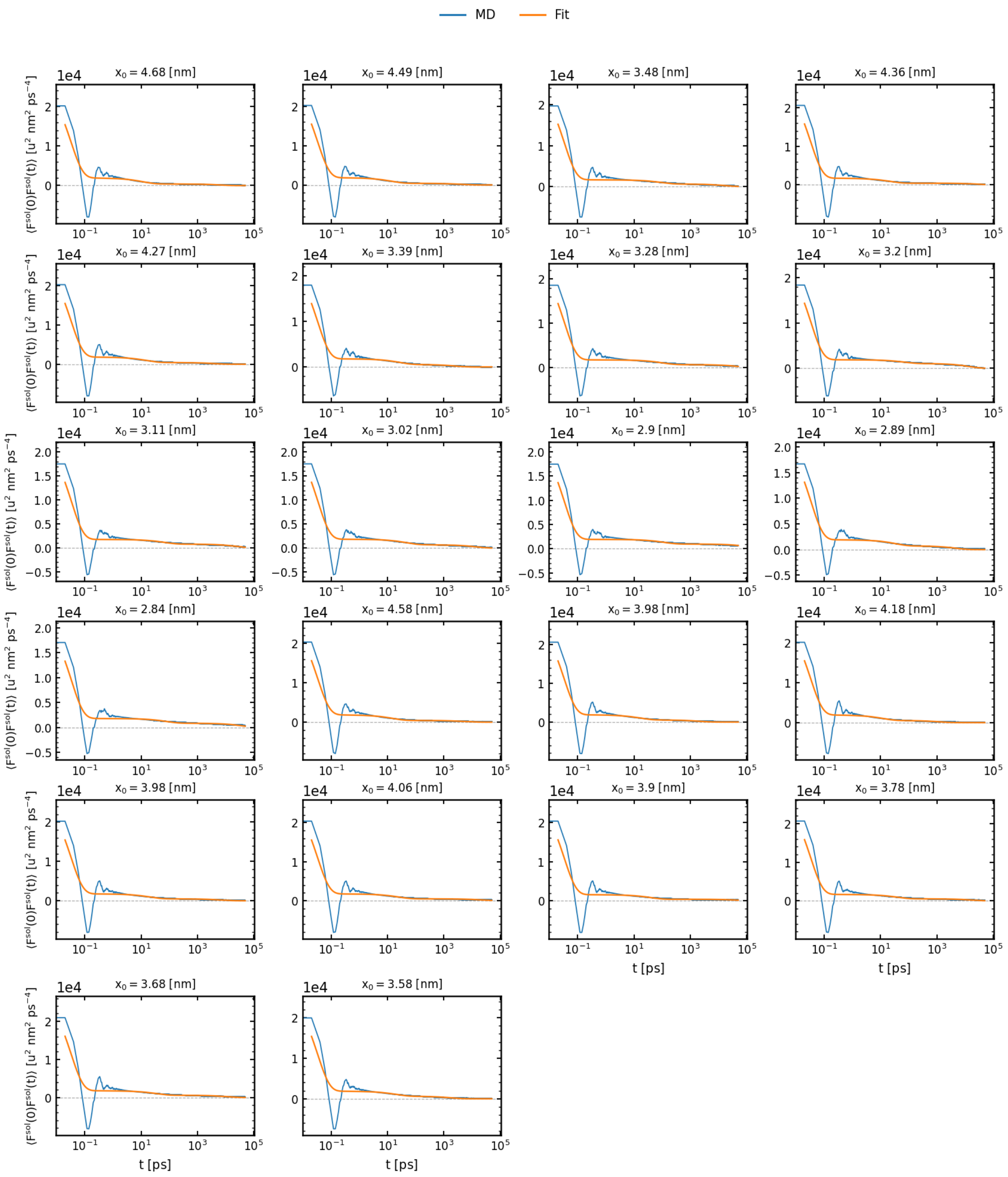


**Supplementary figure 15. Solvent-force autocorrelation functions along the c–NBS coordinate.** Solvent-force autocorrelation functions $\langle F_{sol}(0)F_{sol}(t)\rangle$ computed along the c–NBS (C416 – L458) collective variable for each umbrella sampling window. Blue curves show the molecular dynamics data, and orange curves show the fitted correlation functions used to estimate the solvent-force friction. The fits capture the overall decay of the force correlations along the c–NBS coordinate, while short-time oscillations and long-time noise are not fully resolved by the smooth fitting function. The titles denote $x_0$, the centers of the restraining potentials along the C–NBS collective variable.

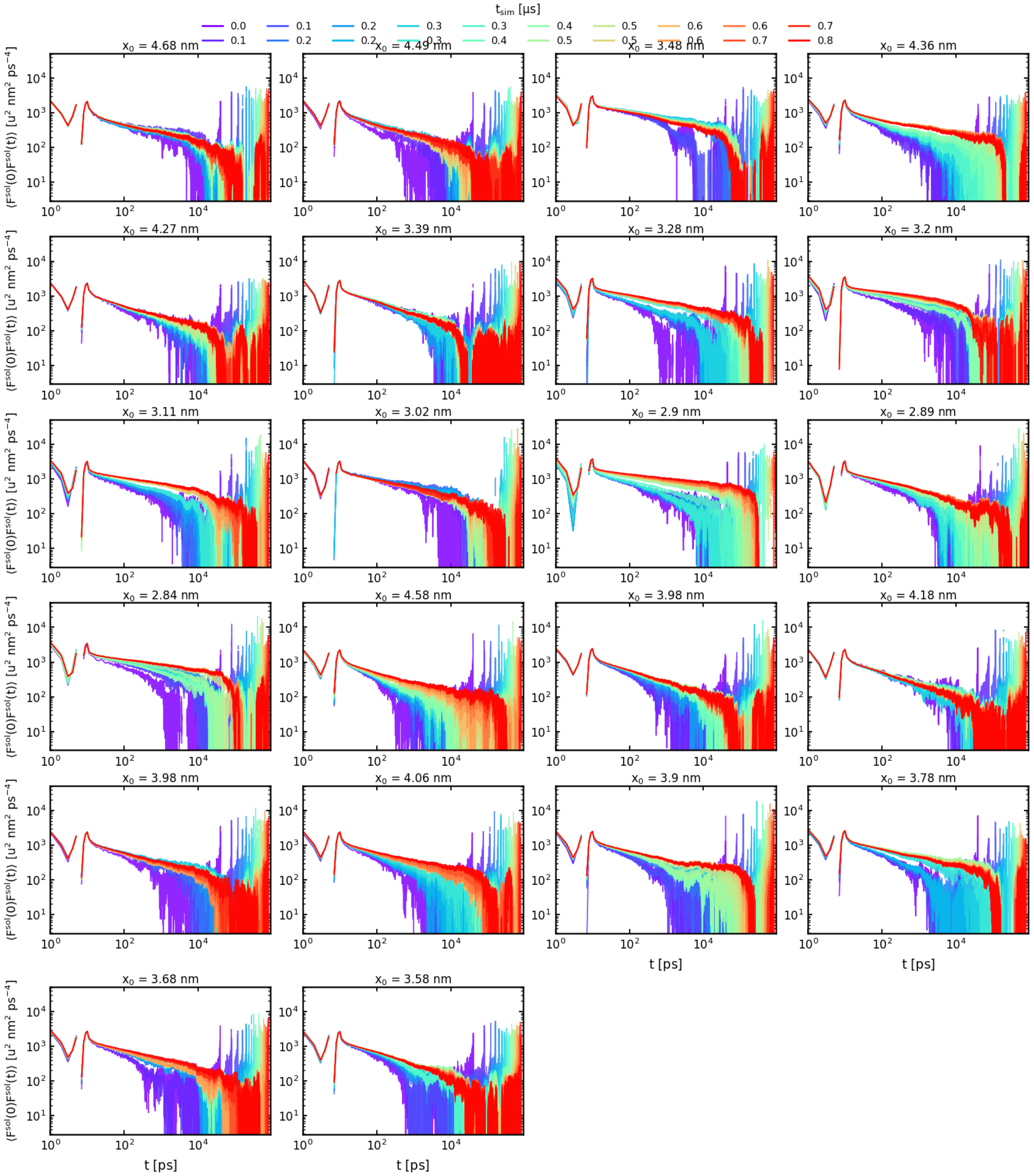


**Supplementary figure 16. Convergence of solvent-force autocorrelation functions along the c–NBS coordinate.** Solvent-force autocorrelation functions $\langle F_{\mathrm{sol}}(0)F_{\mathrm{sol}}(t)\rangle$ along the c–NBS (C416 – L458) collective variable, computed using increasing amounts of simulation data. Colors indicate the cumulative simulation time included in the analysis. The short-time decay is largely reproducible, whereas the long-time tails show substantial statistical fluctuations, as expected for force autocorrelation functions. These data are therefore used primarily to obtain a rough comparison friction estimate rather than a precise window-by-window friction profile along the c–NBS coordinate.

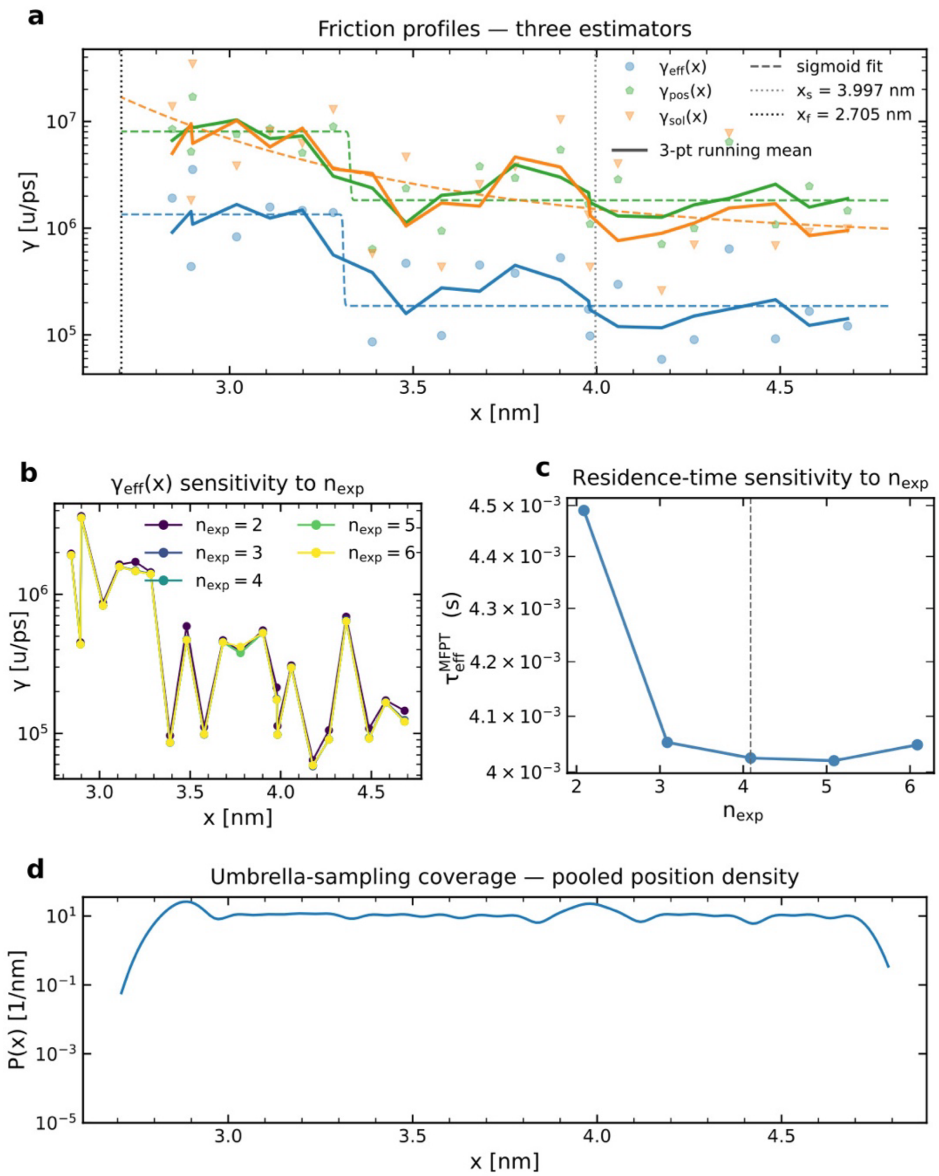


**Supplementary figure 17. (a) Position-dependent friction profiles along the c–NBS collective variable, defined by the C416–L458 distance**. Three correlation-based estimators are shown: the memory-corrected effective friction γeff(x) from the squared position autocorrelation (blue circles), the position-autocorrelation estimate γpos(x) (green pentagons), and the solvent-force estimate γsol(x) (orange triangles). The red curve shows the fitted effective-friction profile γefffit(x) used as input for the MFPT calculation. Vertical dotted lines mark the initial and final c–NBS coordinate values used in the MFPT estimate. Although the individual window estimates scatter substantially, the three estimators give a consistent qualitative ordering, with γpos(x) and γsol(x) generally larger than γeff(x). **(b)** Sensitivity of the fitted γeff(x) profile to the number of exponential components used in the multi-exponential fits of Cxx(t). The near-overlap of the curves indicates that the extracted friction profile is insensitive to the precise choice of nexp over the tested range. **(c)** Corresponding MFPT as a function of the absorbing-boundary position along the c–NBS coordinate. The vertical dashed line marks the final coordinate value used to report the residence/transition time. The MFPT rises across the high-friction/barrier region and reaches a plateau near the final boundary, indicating that the selected endpoint captures the dominant kinetic contribution. **(d)** Sampling coverage along the c–NBS coordinate from the combined umbrella-sampling windows. The broad plateau indicates continuous coverage of the sampled coordinate range, while the drop at the boundaries reflects the edges of the umbrella array. Together, the panels show that the MFPT estimate is based on a continuous sampled coordinate, a stable effective-friction fit, and a friction hierarchy that is robust despite window-to-window scatter.

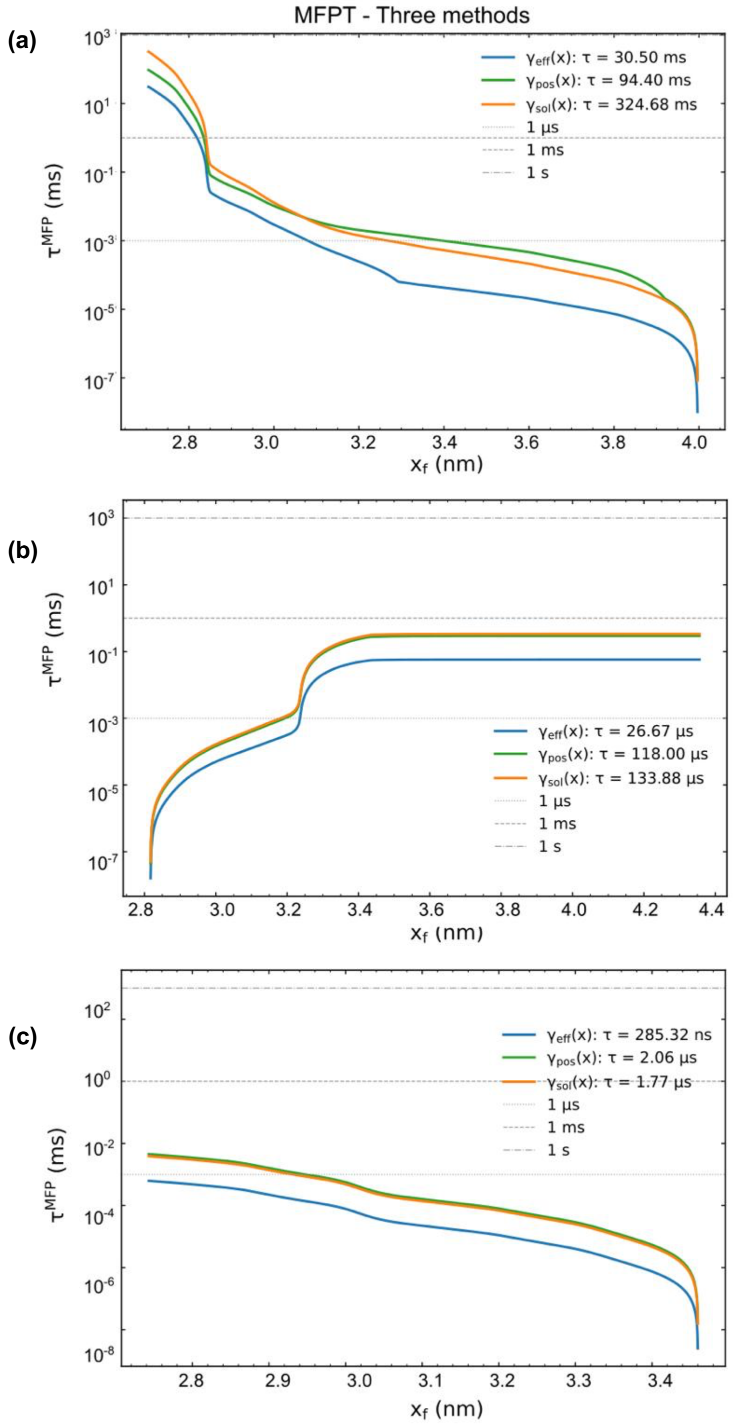


**Supplementary figure 18. MFPT estimates along the a) c–NBS, b) TMD, and c) d-NBS collective variables.** Mean first-passage time $\tau_{MFPT}$ as a function of the absorbing boundary position $x_f$ along the collective variables, computed using three different position-dependent friction profiles. Blue, green, and orange curves correspond to $\gamma_{eff}(x)$, $\gamma_{pos}(x)$, and $\gamma_{sol}(x)$, respectively. Horizontal dashed lines indicate reference times of 1 µs, 1 ms, and 1 s. The spread between the three estimates reflects uncertainty in the friction profile and is used as an approximate uncertainty range for the MFPT. Despite quantitative variation between friction estimators, all estimates place the transition in the millisecond regime.

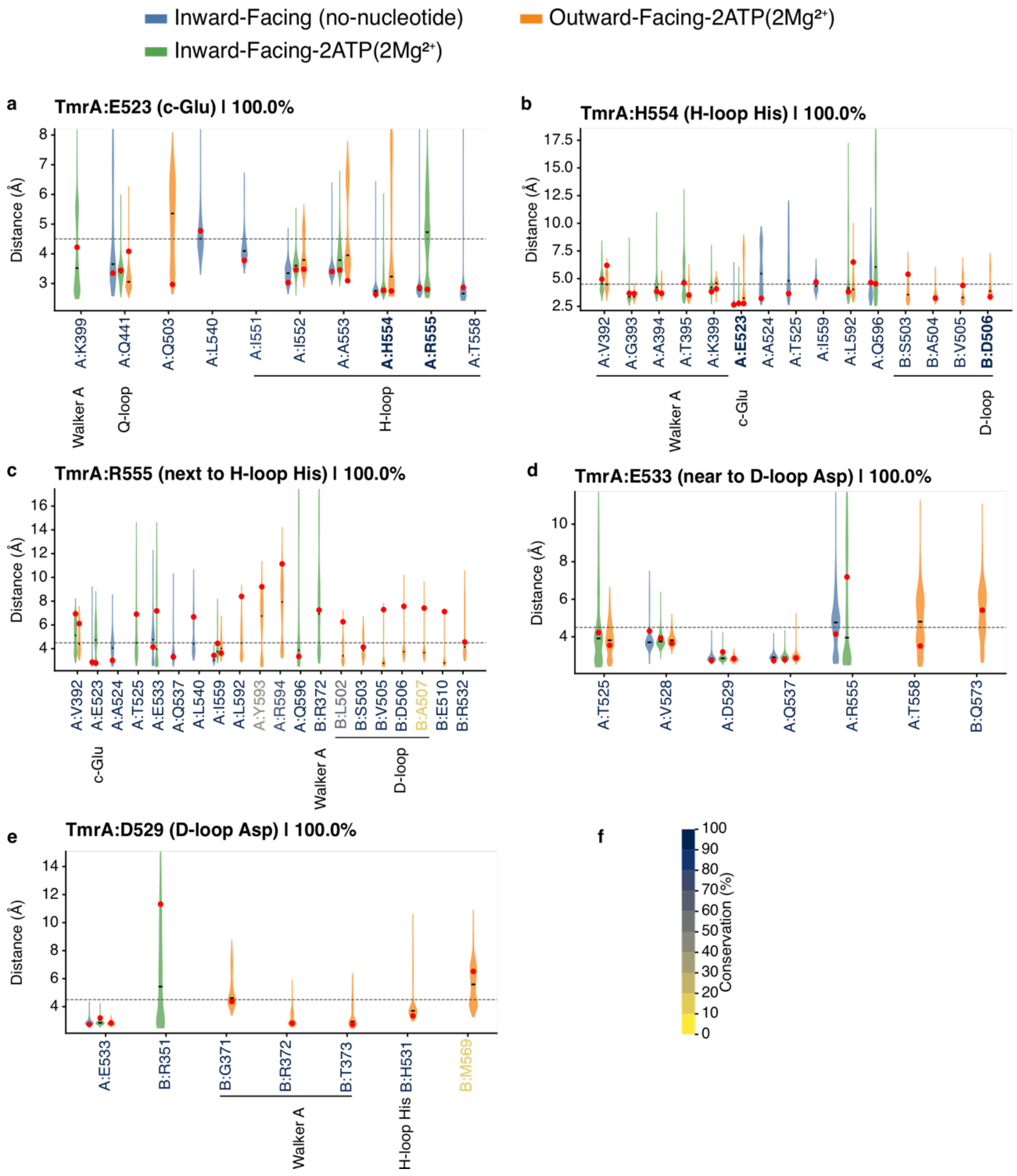


**Supplementary figure 19. Comparative contact distributions for key residues forming IF-lock and OF-lock in TmrA.** The plots correspond to the IF-lock, IF-transition lock, and the OF-lock as presented in Figure 3a-c. The violin plots show minimum heavy-atom distance distributions between the key residues (**a**) TmrA-E523, (**b**) TmrA-H554, (**c**) TmrA-R555, TmrA-D529, (**d**) TmrA-E533, (**e**) TmrA-D529 (which are 100% conserved), and their contacting partner residues (including those in TmrB) across three conformational states: inward-facing (apo / non-nucleotide), inward-facing-ATP-$Mg^{2+}$, and outward-facing-ATP-$Mg^{2+}$. Each condition was sampled using 10 x 500 ns MD simulations. Violin plot colors indicate the simulation condition, and red dots mark the corresponding starting distances from the initial structure of each condition. The dashed horizontal line denotes the 4.5 Å contact cutoff, beyond which the interaction is too weak. Only partner residues with contact occupancy greater than 30% in at least one condition are shown. Partner-residue labels are colored according to sequence conservation as indicated by the conservation color scale (**f**), and the key residues presented are fully conserved (indicated with 100% mark, as shown in Supplementary figure 22). Distances were calculated as the minimum heavy-atom separation between residue pairs, including both backbone and sidechain atoms but excluding hydrogens. Candidate partner residues were prefiltered independently for each condition using the corresponding starting structure, retaining residues within 15 Å of each anchor residue before trajectory-wide distance analysis. Residue pairs within 3 sequence positions in the same chain were excluded, whereas inter-chain residue pairs were retained.

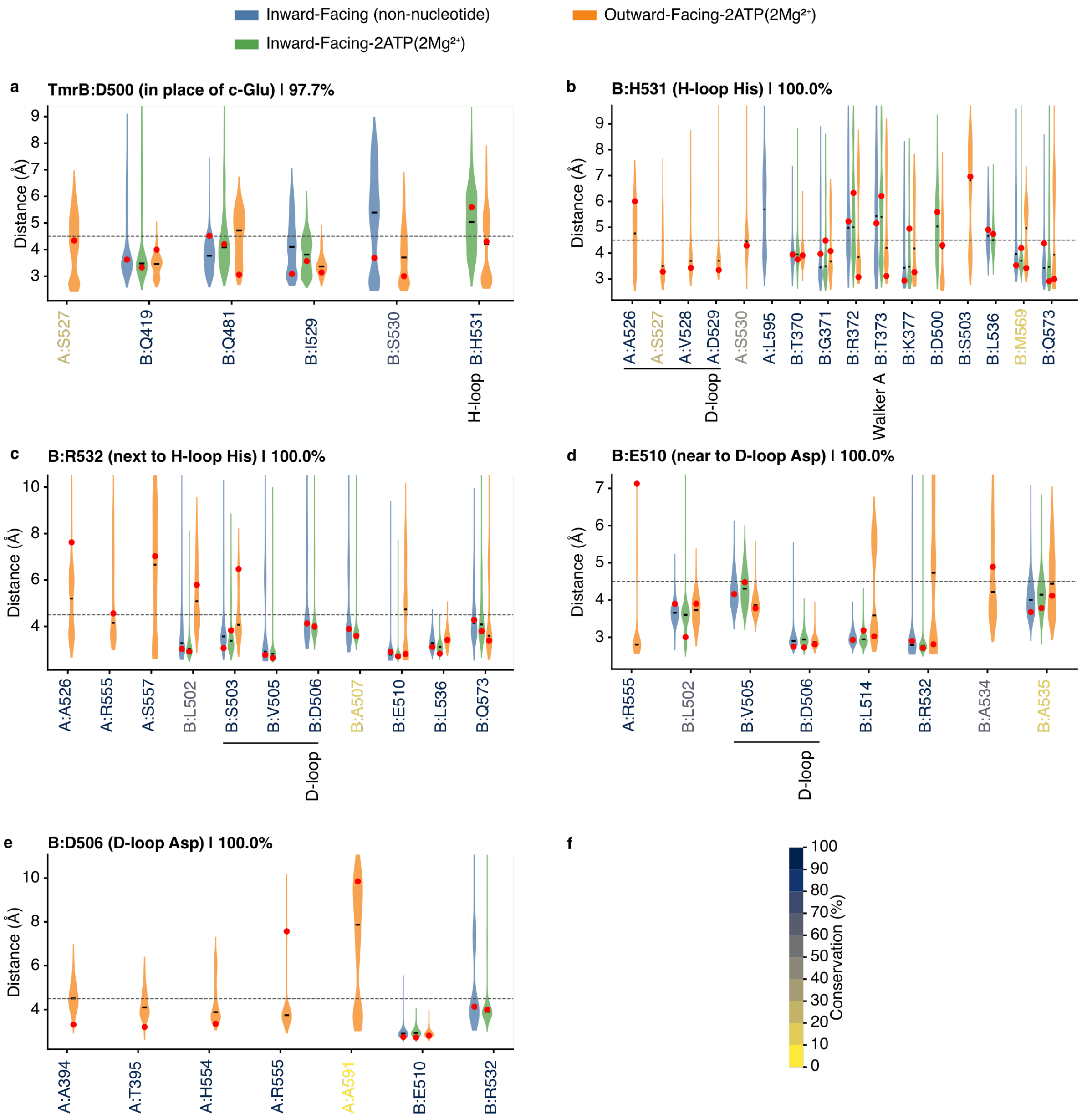


**Supplementary figure 20. Comparative contact distributions for key residues forming IF-lock and OF-lock in TmrB.** The plots correspond to the IF-lock, IF-transition lock, and the OF-lock as presented in Figure 3a-c. Violin plots show minimum heavy-atom distance distributions between the key residues TmrB-D500, TmrB-D506, TmrB-E510, TmrB-H531, and TmrB-R532 and their contacting partner residues across three conformational states: inward-facing (non-nucleotide), inward-facing-ATP-$Mg^{2+}$, and outward-facing- ATP-$Mg^{2+}$. Each condition was sampled using 10 x 500 ns MD simulations. Violin colors indicate the simulation condition, and red dots mark the corresponding starting distances from the initial structure of each condition. The dashed horizontal line denotes the 4.5 Å contact cutoff, beyond which the interaction is too weak. Only partner residues with contact occupancy greater than 30% in at least one condition are shown. Partner-residue labels are colored according to sequence conservation as indicated by the conservation color scale (f), and the key residues presented are fully conserved (indicated with 100% mark, as shown in Supplementary figure S20). Distances were calculated as the minimum heavy-atom separation between residue pairs, including both backbone and sidechain atoms but excluding hydrogens. Candidate partner residues were prefiltered independently for each condition using the corresponding starting structure, retaining residues within 15 Å of each anchor residue before trajectory-wide distance analysis. Residue pairs within 3 sequence positions in the same chain were excluded, whereas inter-chain residue pairs were retained.

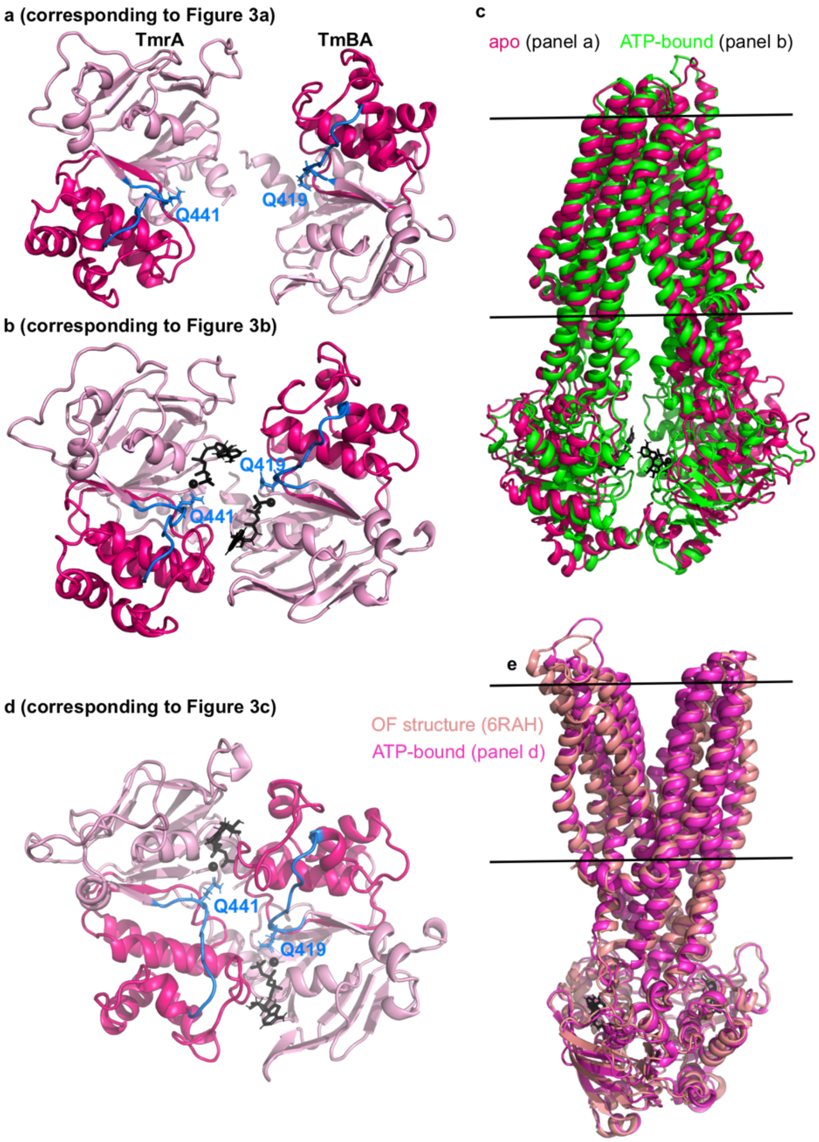


**Supplementary figure 21**. Snapshots from post-equilibration MD simulations for the apo, inward-facing-ATP-$Mg^{2+}$, and outward-facing-ATP-$Mg^{2+}$ structures. The corresponding contact distributions are presented in Supplementary figures 19-20. The images show the global structure of the NBSs and the TMDs corresponding to the IF-lock, IF-transition lock, and the OF-lock as presented in Figure 3a-c. (**a**) The apo, (**b**) $Mg^{2+}$-ATP-bound inward-facing, and (**d**) $Mg^{2+}$-ATP-bound outward-facing TmrAB snapshots from MD simulations are presented. An overlay of the respective MD snapshots is shown (**c**) with the apo structure (PDB ID: 6RAG) or (**e**) the OF structure (PDB ID: 6RAH). The Q-loop is highlighted in blue, and the conserved glutamines are shown in stick representation. The $Mg^{2+}$ ions (as black spheres) and ATPs (in black stick representation) are highlighted.

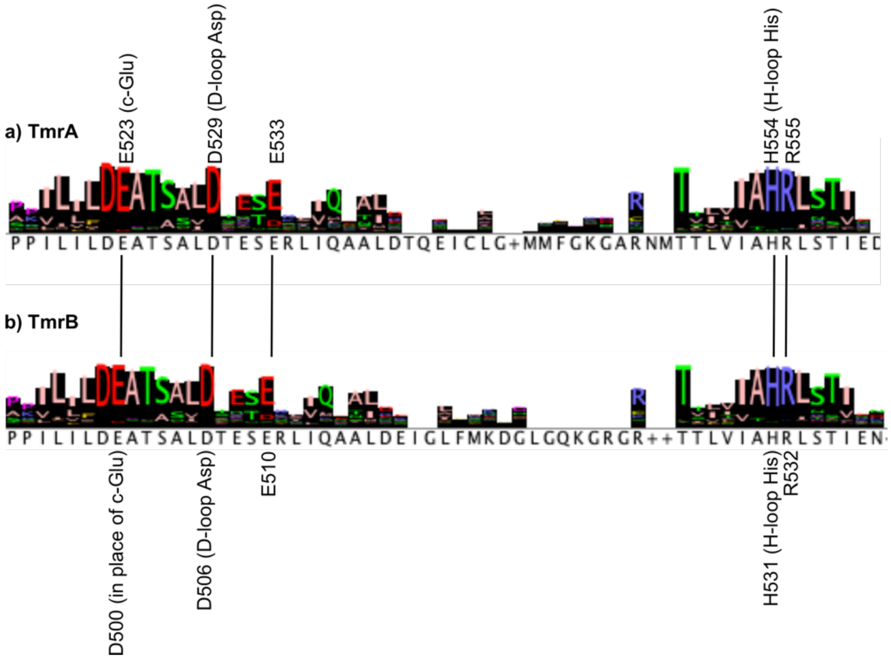


**Supplementary figure 22**. **Global conservation analysis for the key residues constituting the IF-lock and OF-lock in TmrA and TmrB.** The key residues constituting the IF-lock and OF-lock are indicated. The only missing residue is Walker A Ser/Thr, (which is rarely an Asn, at the third position in the Walker A sequence GXTXXGK[S/T]), which is conserved across ABC proteins. Five hundred sequences from the ABC superfamily matching closest with TmrA and TmrB were used for the analysis. The catalytic glutamate (c-Glu), D-loop Asp, and H-loop His are invariant among ABC proteins. The analysis revealed that the Glu (or rarely an Asp) right next to the D-loop Asp (TmrA E533, TmrB E510) and the Arg right next to the H-loop His (TmrA R555 and TmrB R532), which are part of the molecular locks, are conserved in type IV and type VI ABC exporters, and the distantly related Rad50 protein.

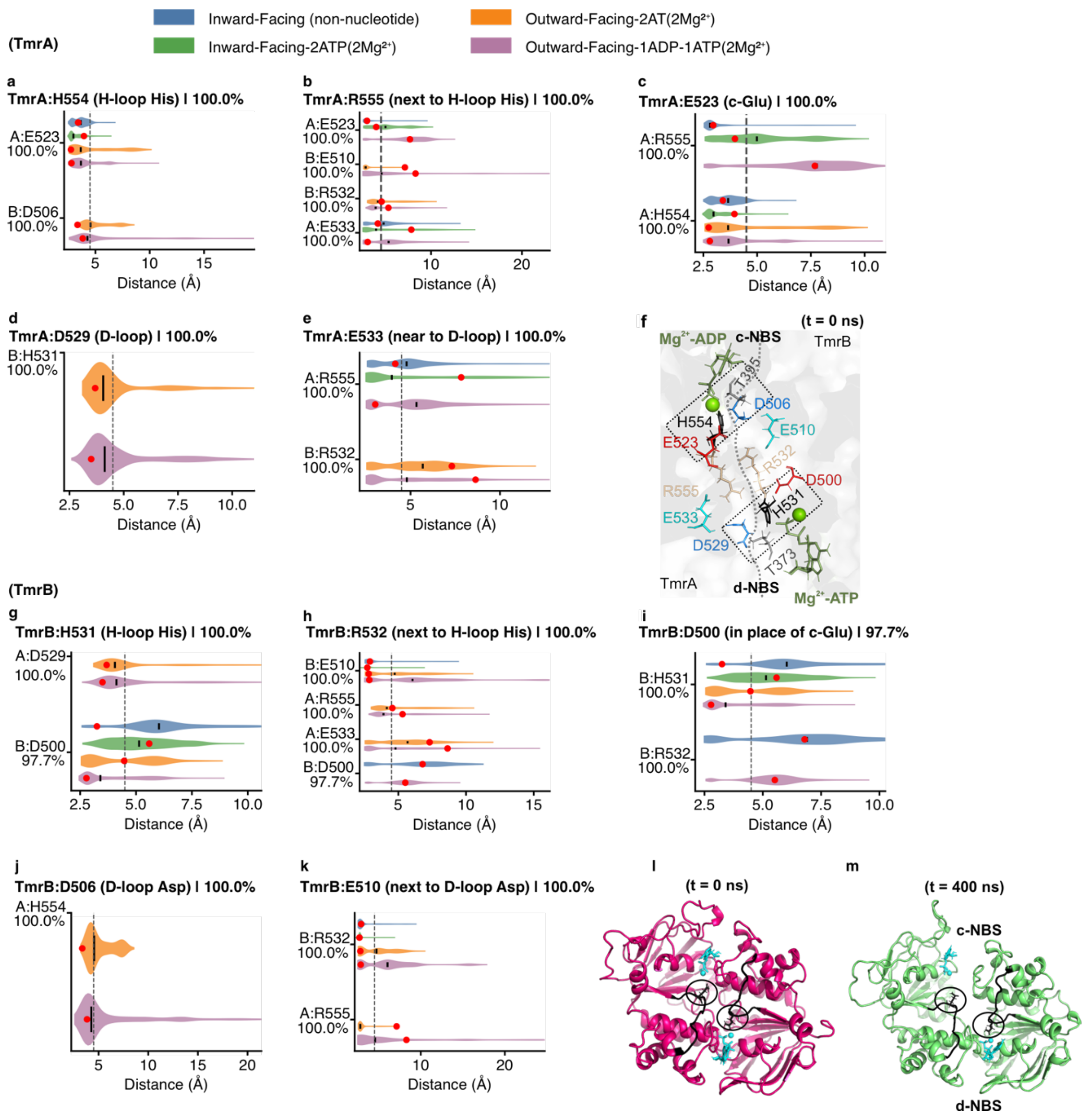


**Supplementary figure 23. Contact distribution for the key residues forming the IF-lock and the OF-lock in wild-type TmrAB during reverse (OF→IF) transition.** The violin plots represent minimum heavy-atom distance distributions for the key residues with their partners forming the molecular locks (panel **f**, also see Figure 3a-c) during transition from ADP-$Mg^{2+}$ (c-NBS) + ATP-$Mg^{2+}$ bound OF structure towards the IF conformation. The figure provides a more detailed view of the changes during reverse transition as presented in Figure 4. For a direct comparison, contact distributions in the inward-facing (non-nucleotide), inward-facing-ATP-$Mg^{2+}$, and outward-facing- ATP-$Mg^{2+}$ states, which are presented in Supplementary figures 19-20, are overlaid. Each state was sampled using 10 x 500 ns MD simulations. Violin colors denote the simulation condition, red dots indicate the corresponding starting distances, and the dashed vertical line marks the 4.5 Å contact cutoff. Only partner residues exhibiting contact occupancy greater than 10% in at least one condition are included. Partner-residue conservation values are shown beneath the y-axis labels, while key/anchor-residue conservation is given in each panel title. Distances were defined using the minimum heavy-atom separation, including backbone and sidechain atoms but excluding hydrogens. Residue pairs within 3 sequence positions in the same chain were excluded; inter-chain residue pairs were retained. In the last panels (**l**-**m**), the orientation of the Q-loops (in black) during reverse transition in the ADP-$Mg^{2+}$ (c-NBS) + ATP-$Mg^{2+}$ (d-NBS) structure is shown. The bound $Mg^{2+}$ (as a sphere) and the nucleotides (in stick representation) are highlighted in cyan.

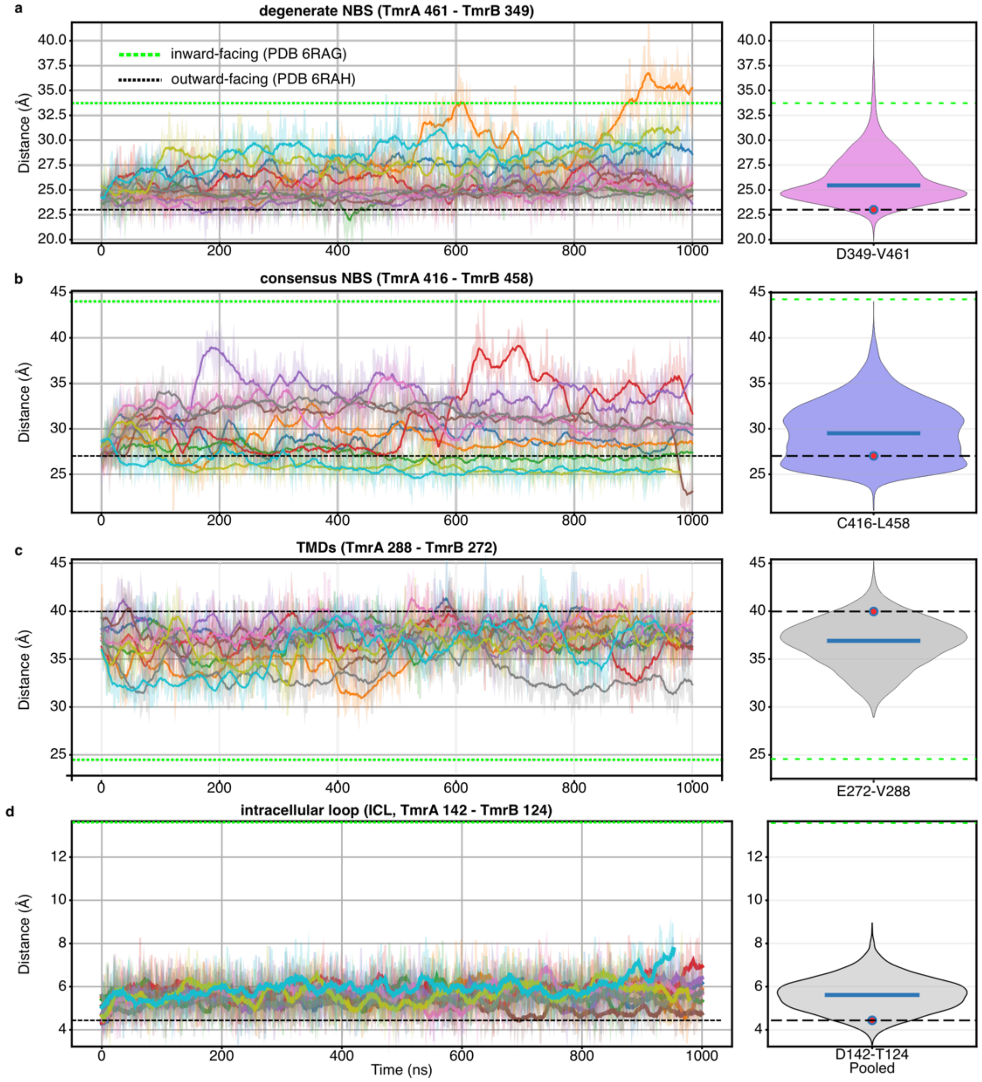


**Supplementary figure 24**. **Time evolution and pooled distance distributions in the outward-facing ADP-$Mg^{2+}$ (c-NBS) + ATP–$Mg^{2+}$ (d-NBS)-bound TmrAB (WT) during reverse transition.** The trajectories correspond to the selected violin plots presented in Figure 4a-c, and further interactions are presented in Supplementary figure 23. The distances monitored are (**a**) V461-D349 (d-NBS), (**b**) C416–L458 (c-NBS), (**c**) V288–E272 (TMDs), and (**d**) D142–T124 (ICL). In the left panels, each colored trace represents one independent trajectory (10 x 500 ns runs), with faint lines showing the raw data and bold lines showing the smoothed trajectories. In the right panels, violin plots show the pooled distribution of each distance across all trajectories; the blue horizontal line marks the median, and the red circle marks the mean value at the starting frame. For structural comparison, the outward-facing reference structure is indicated by a black dashed line and the inward-facing reference structure by a green dotted line in both the time-series and violin panels. Together, these comparisons show how the simulated outward-facing ensemble evolves towards inward-facing conformations.

**Supplementary Table 1. Kinetic analysis of the TMDs (TmrA 288 – TmrB 272) in the presence of ATP (EDTA).** Values for the fitted parameters with standard error (SE) are shown (parameters correspond to the fitting of data presented in Figure 1e, see methods for further details).

| T (K) | $P_{eq}$ | SE ($P_{eq}$) | $k_{tot}$ (×10$^{-3}$) (S$^{-1}$) | SE ($k_{tot}$) (×10$^{-3}$) (S$^{-1}$) | $k_1$ (×10$^{-3}$) (S$^{-1}$) | SE ($k_1$) (×10$^{-3}$) (S$^{-1}$) | $k_{-1}$ (×10$^{-3}$) (S$^{-1}$) | SE ($k_{-1}$) (×10$^{-3}$) (S$^{-1}$) | $K_{eq}$ | SE ($K_{eq}$) |
|---|---|---|---|---|---|---|---|---|---|---|
| 303.15 | 0.32 | 0.02 | 5.34 | 1.35 | 1.68 | 0.35 | 3.66 | 1.01 | 0.46 | 0.04 |
| 313.15 | 0.48 | 0.02 | 7.79 | 1.19 | 3.76 | 0.47 | 4.03 | 0.74 | 0.93 | 0.07 |
| 323.15 | 0.56 | 0.03 | 21.01 | 4.75 | 11.70 | 2.35 | 9.30 | 2.49 | 1.26 | 0.14 |
| 333.15 | 0.60 | 0.02 | 57.73 | 16.22 | 34.47 | 9.33 | 23.26 | 7.09 | 1.48 | 0.15 |
| 338.15 | 0.64 | 0.03 | 62.26 | 12.79 | 39.99 | 7.44 | 22.27 | 5.63 | 1.80 | 0.20 |

**Supplementary Table 2. Kinetic analysis of the c-NBS (TmrA 416 – TmrB 458) in the presence of ATP (EDTA).** Values for the fitted parameters with standard error (SE) are shown (parameters correspond to the fitting of data presented in Figure 1f, see methods for further details).

| T (K) | $P_{eq}$ | SE ($P_{eq}$) | $k_{tot}$ (×10$^{-3}$) (S$^{-1}$) | SE ($k_{tot}$) (×10$^{-3}$) (S$^{-1}$) | $k_1$ (×10$^{-3}$) (S$^{-1}$) | SE ($k_1$) (×10$^{-3}$) (S$^{-1}$) | $k_{-1}$ (×10$^{-3}$) (S$^{-1}$) | SE ($k_{-1}$) (×10$^{-3}$) (S$^{-1}$) | $K_{eq}$ | SE ($K_{eq}$) |
|---|---|---|---|---|---|---|---|---|---|---|
| 303.15 | 0.38 | 0.07 | 1.78 | 0.68 | 0.68 | 0.13 | 1.10 | 0.56 | 0.62 | 0.20 |
| 313.15 | 0.42 | 0.02 | 10.31 | 1.51 | 4.36 | 0.50 | 5.95 | 1.03 | 0.73 | 0.05 |
| 323.15 | 0.50 | 0.02 | 19.29 | 2.90 | 9.73 | 1.23 | 9.57 | 1.71 | 1.02 | 0.08 |
| 333.15 | 0.48 | 0.01 | 46.98 | 4.17 | 22.44 | 1.80 | 24.54 | 2.42 | 0.91 | 0.03 |

**Supplementary Table 3. Summary of the molecular dynamics (MD) simulations performed in this study.**

| Setup ID | Purpose / figure linkage | Starting structure and state | Protein / nucleotide condition | Replicas or windows | Length | Total sampling to report | Temperature / pressure | Count as independent new sampling? |
|---|---|---|---|---|---|---|---|---|
| MD-01 | Unbiased forward IF to OCC production MD; Main Fig. 2a; Supplementary Figs. 5-II and 5-III | Inward-facing TmrAB; $Mg^{2+}$-ATP-bound state | WT TmrAB with bound $Mg^{2+}$-ATP | 10 independent replicas | 500 ns each | 5.0 µs | 333.15 K / 60 °C; 1 bar; POPE/POPG (3:1) | Yes |
| MD-02 | Unbiased forward IF to OCC production MD; Main Fig. 2a; Supplementary Figs. 5-I and 5-III | Inward-facing TmrAB; $Mg^{2+}$-ATP-bound state | E-to-Q variant with bound $Mg^{2+}$-ATP | 10 independent replicas | 500 ns each | 5.0 µs | 333.15 K / 60 °C; 1 bar; POPE/POPG (3:1) | Yes |
| MD-03 | OCC to OF production MD; Main Fig. 2a text; Supplementary Fig. 6 | Occluded TmrAB; $Mg^{2+}$-ATP-bound state | E-to-Q variant with bound $Mg^{2+}$-ATP | 6 independent replicas | 1.7 µs each | 10.2 µs | 333.15 K / 60 °C; 1 bar | Yes |
| MD-04 | Lock-state contact-distribution simulations/analysis; Main Fig. 3a-c; Supplementary Figs. 17-19 | Inward-facing TmrAB; IF apo/non-nucleotide | WT TmrAB lock-state systems unless variant is explicitly stated in the figure legend | 5 Independent replicas | 500 ns each | 2.5 µs per condition | 333.15 K / 60 °C; 1 bar; same production MD conditions | yes |
| MD-05 | Targeted MD before umbrella sampling for PMF/MFPT; Main Fig. 2e-l; Supplementary Figs. 8-I, 8-II, 9-16 | WT $Mg^{2+}$-ATP-bound IF structure, moved toward OCC and OF conformations | WT TmrAB with $Mg^{2+}$-ATP | no | 400 ns per window | 0.8 µs | 310 K; umbrella spring constant 1000 kJ $mol^{-1}$ $nm^{-2}$ | Yes |
| MD-06 | Reverse-transition perturbation MD after γ-phosphate/Pi removal; Main Fig. 4; Supplementary Figs. 21-22 | Outward-facing $Mg^{2+}$-ATP-bound state after γ-phosphate/Pi removal at the c-NBS | WT TmrAB; $Mg^{2+}$-ADP/$Mg^{2+}$-ATP post-perturbation state after removal at c-NBS | 10 independent replicas | 500 ns each | 5.0 µs | 333.15 K / 60 °C; 1 bar; POPE/POPG (3:1) | Yes |
| MD-07 | Reverse-transition controls; Main Fig. 4a-c,d-g | Outward-facing controls | WT without γ-phosphate removal; E-to-Q with γ-phosphate/Pi removal at c-NBS | 10 independent replicas | 500 ns each | 5.0 µs | 333.15 K / 60 °C; 1 bar; POPE/POPG (3:1) | Yes |
